# Environmental fluctuations alter the competitive trade-offs of group size in a social primate

**DOI:** 10.1101/2025.08.12.669921

**Authors:** Odd T. Jacobson, Margaret C. Crofoot, Genevieve E. Finerty, Brendan J. Barrett, Susan E. Perry

## Abstract

Larger animal groups are widely understood to require more space and expend more energy to mitigate the foraging costs of within-group competition. Yet between-group interactions and shifting resource distributions can obscure links between group size and behavior, making responses to demographic change difficult to predict. Using 33 years of observational data from 12 neighboring white-faced capuchin (*Cebus imitator*) groups in Costa Rica, combined with remotely sensed environmental data, we show that within- and between-group competition jointly shape space use, with their relative importance shifting with seasonal and interannual climate cycles. Larger groups compensated for reduced per-capita foraging efficiency by expanding into less-exploited areas over longer timescales rather than increasing daily travel. Notably, this expansion disproportionately encroached on smaller neighboring groups. In the dry season, resource confinement to riparian zones increased intergroup encounters and reduced overlap, with larger groups occupying the highest-quality areas. Climatic extremes linked to El Niño and La Niña exacerbated within-group foraging costs for large groups, whereas intermediate anomalies relaxed these constraints and amplified the benefits of between-group competitive ability. Our findings show that environmental variation shifts the trade-offs of within- and between-group competition, shaping how group-living animals adjust to changing social and ecological conditions.

## 1 Main

Functional traits of animals (e.g., body size, life-history strategy, social structure) carry both costs and benefits that depend on specific ecological conditions [1, 2]. Understanding how environmental change alters the balance of these trade-offs is key to explaining their wide variation across and within species (and populations), as well as to forecasting evolutionary trajectories [3, 4]. Group size (an emergent demographic trait) represents one such trade-off for group-living animals, as it affects both within- and between-group resource competition, but often in opposing ways [5–7].

Within-group competition imposes a cost by reducing individual foraging efficiency, thereby setting an upper limit on group size (i.e., the Ecological Constraints Model of Group Size; [8–10]), which primarily reflects scramble (indirect) rather than contest (direct) competition among group members [5]. Consequently, larger groups face higher energetic costs from increased resource requirements and travel distances, driven by greater collective metabolic demands and faster resource depletion [11, 12]. They also often require larger home ranges (e.g., [13–16]; but see [17])—similar to how individual space use scales allometrically with body size [18].

While larger groups typically face greater costs from within-group competition, they often hold a competitive edge in between-group contests, leveraging their numerical advantage to exclude smaller neighbors from key resources [17, 19–22]. This advantage is especially pronounced when resources, such as fruiting trees, are distributed in defensible patches that vary in quality, allowing larger groups to monopolize the most valuable patches [6]. Larger groups also benefit indirectly, as smaller groups may avoid high-risk areas—granting larger groups access to a wider array of food patches with reduced opposition [23, 24]. With more (and better) foraging options available, larger groups can access richer and less-depleted patches, potentially mitigating the costs of within-group competition and contributing to higher survival and reproductive success [6, 25–27]. Ultimately, group size reflects trade-offs between the costs (e.g., within-group competition, disease transmission) and benefits (e.g., between-group competitive advantage, reduced predation risk) of group living [28, 29]. Still, group sizes often vary widely within populations and fluctuate temporally, suggesting that this balance is mediated by additional ecological and social factors [30–32]. Demographic turnover reshapes the distribution of group sizes across the population, altering both ecological constraints and the balance of power among neighboring groups [22, 33]. Simultaneously, local heterogeneity and shifting environmental conditions influence both the intensity of within-group competition and the cost-effectiveness of resource defense [34–36]. Smaller groups may benefit when resources are abundant or widely distributed, making them less practical to defend— thereby reducing the importance of intergroup dominance [19]. In contrast, during resource-scarce periods, larger groups may gain an advantage by using their numerical superiority to outcompete smaller groups for remaining food patches [27, 37]. This advantage, however, could be offset if larger groups incur disproportionately high costs from within-group competition under the same conditions [38]. Thus, which group sizes are favored is likely context-dependent, shifting with demographic and environmental change [4,28, 39].

The Ecological Constraints Model is a widely accepted framework for understanding how within-group competition limits group size in social animals [8, 9, 11, 40–43]. As with any broad framework, however, real-world ecological complexity introduces important nuance. The interdependence of within- and between-group competition, along with the dynamic nature of landscapes and competitive interactions, often obscures causal links between group size and behavior. Thus, how costs of within-group competition interact with benefits of between-group competition at the population level—particularly as resource availability fluctuates— remains unresolved. Clarifying how this interaction unfolds over time is critical for understanding the evolution of sociality, the ecological conditions under which it is favored, and its trajectory in a rapidly changing world. Such insight requires long-term datasets from multiple, neighboring social groups that encompass meaningful demographic and ecological variation— data that are logistically difficult to obtain. Most previous research has therefore examined within- and between-group competition in isolation (often cross-sectionally), lacking either the temporal depth or number of social groups needed to explore their interaction.

Drawing upon 33 years of data on 12 white-faced capuchin monkey groups from the Lomas Barbudal Monkey Project in Costa Rica (10*^◦^*29–32’N, 85*^◦^*21–24’W), we examine how group size and environmental variation structure population-level patterns of resource competition in a group-living animal [44]. Lomas Barbudal lies within one of the last remaining fragments of tropical dry forest [45]—a biome characterized by a highly seasonal climate with a distinct wet season (May–November) and a harsh dry season (December–April) (Figure 1b; [46]). Seasonal shifts dramatically alter resource distribution and availability. Most of the forest is deciduous; during the dry season water, shade, and food become concentrated in evergreen riparian zones (subsection S1.3). The region is also highly sensitive to interannual climatic fluctuations associated with the El Niño–Southern Oscillation (ENSO; El Niño and La Niña; subsubsection S1.3.2; [47]). In the tropical Americas, El Niño events typically produce abnormally hot, dry conditions, whereas La Niña events bring abnormally cool, wet conditions [48].

**Figure 1:**
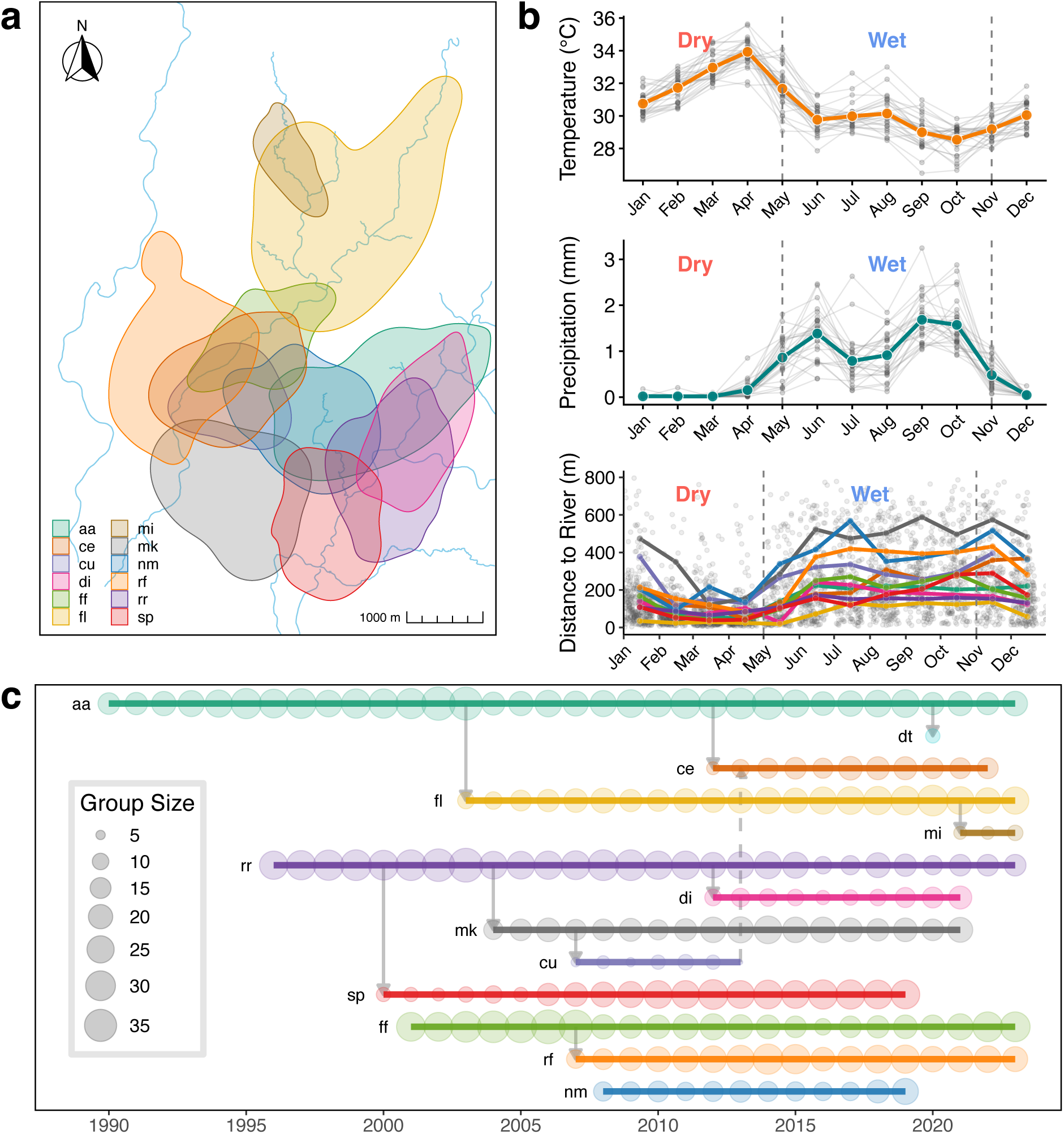
Environmental and demographic context. a: Annual home ranges of 12 white-faced capuchin study groups (95% auto-correlated kernel density contours) at the Lomas Barbudal Monkey Project, Costa Rica. Colors correspond to group identity. Estimates are derived from handheld GPS-tracking data collected in 2012, except force and di (formed after fissions in late 2012; shown using 2013 data) and mi (formed after a 2021 fission; shown using 2021 data). Group dt is not shown due to insufficient location data. b: Weather data near the Lomas Barbudal Biological Reserve from the ERA5 reanalysis dataset (1990–1993, 1997–1999, 2001–2020). Top panel shows temperature (*^◦^*C); middle panel shows total precipitation (mm); bottom panel shows each study group’s mean distance to river. For temperature and precipitation, grey lines and points show monthly means per year; colored lines and points (orange for temperature, blue-green for precipitation) show the monthly mean across years. For distance to river, grey points represent mean daily distance to the river; colored points and lines show monthly means by group. Dashed vertical lines mark typical transitions between dry and wet seasons. c: Time series of data collection for all 13 study groups from 1990 to 2023. Group abbreviations indicate monitoring start dates or formation via permanent fission. Arrows denote fission events (solid) and a single fusion event (dashed).

This system offers a rare opportunity to study how long-term demographic and environmental fluctuations shape competition and space-use dynamics, due to both the exceptional breadth of the dataset and the pronounced environmental variation in the region. Capuchins in this population live in cohesive multi-male, multi-female groups of 5–40 individuals (mean = 18.8; [44]) that persist for years or decades, although occasionally split permanently into two independent units with distinct home ranges (nine events over the study period; Figure 1c). Capuchins are primarily frugivorous (∼50-55%), with invertebrates comprising most of the remaining percentage (based both on time spent feeding [49] and proportion of food consumed [50]). Groups are range-resident for decades and their home ranges overlap extensively (Figure 1a). Inter-group contests are hostile and occur frequently (∼1 per week per group) [51, 52].

Here, we integrate conventional *group-level* predictions of the Ecological Constraints Model— linking group size to indicators of within-group competition— with *dyadic* comparisons of neighboring groups to test how relative group size influences between-group relationships. First, we evaluate annual and seasonal relationships between group size and (a) fruit foraging efficiency, (b) daily path length, (c) return rate to previously used areas, (d) home range area, and (e) home range quality. Next, we apply hierarchical social relations models to evaluate how relative group sizes within group-dyads predict home range overlap and encounter rates, and how these relationships vary with seasonal shifts in vegetation productivity (see Table 1 for biological interpretation of each response variable). Finally, we investigated how seasonal severity influences within- and between-group competition, with the aim of assessing how climatic anomalies linked to ENSO modify the competitive trade-offs of group size. Together, these analyses provide, to our knowledge, the first longitudinal tests of how group-living animals adjust both within- and between-group competition in response to group size and environmental change (see subsection S1.4 for *a priori* predictions and causal pathways). Our findings highlight the dynamic trade-offs of group living and show how ecological and social pressures jointly shape space-use strategies— factors that ultimately determine access to resources and fitness.

**Table 1:** Summary of response variables.

| Response variable | Abbreviation | Level | Competition type | Interpretation |
| --- | --- | --- | --- | --- |
| Per Capita Fruit Intake Rate | FIR | Individual | Within-group | Individual energy intake from fruit |
| Daily Path Length | DPL | Group | Within-group | Daily energy expenditure |
| Revisitation Rate | RR | Group | Within-group | Frequency of returns to previously used areas |
| Home Range Area | HRA | Group | Within-group | Long-term space requirement |
| Home Range NDVI | HRQ | Group | Between-group | Home range quality |
| Proportional Overlap | $PO_{fn}$ | Group-dyad (asymmetric) | Between-group | Extent of neighbor access to focal range |
| Encounter Rate | ER | Group-dyad (symmetric) | Between-group | Frequency of intergroup encounters |

## 2 Results

### 2.1 Larger groups face foraging costs but do not increase daily travel

To evaluate how group size influences resource competition, we combined long-term movement and observational data with continuous-time movement modeling (ctmm) and Bayesian generalized linear multilevel models (GLMMs) to estimate five response variables: (a) per capita daily fruit intake rate, (b) daily path length, (c) mean revisitation rate across the home range, (d) home range area, and (e) home range quality, assessed via normalized difference vegetation index (NDVI)— a satellite-derived measure of vegetation greenness. All GLMMs were run at both annual and seasonal scales.

We found strong evidence for a negative association between group size and per-capita fruit intake rate, but little to no association with daily path length in either season (Figure 2a). Revisitation rates were considerably lower in larger groups at the annual scale and during the wet season (*β*_annual_ = −0.11[−0.23, 0.00]; *PP >* 0 = 0.07; *β*_wet_ = −0.22[−0.34, −0.09]; *PP >* 0 = 0.008), but showed no relationship in the dry season (*β*_dry_ = −0.06[−0.2, 0.08]; *PP >* 0 = 0.262). Larger groups consistently occupied larger home ranges across both scales, with a stronger effect observed in the wet season (*β*_wet_ = 0.36[0.24, 0.49]; *PP >* 0 = 1; *β*_dry_ = 0.27[0.13, 0.41]; *PP >* 0 = 0.998).

**Figure 2:**
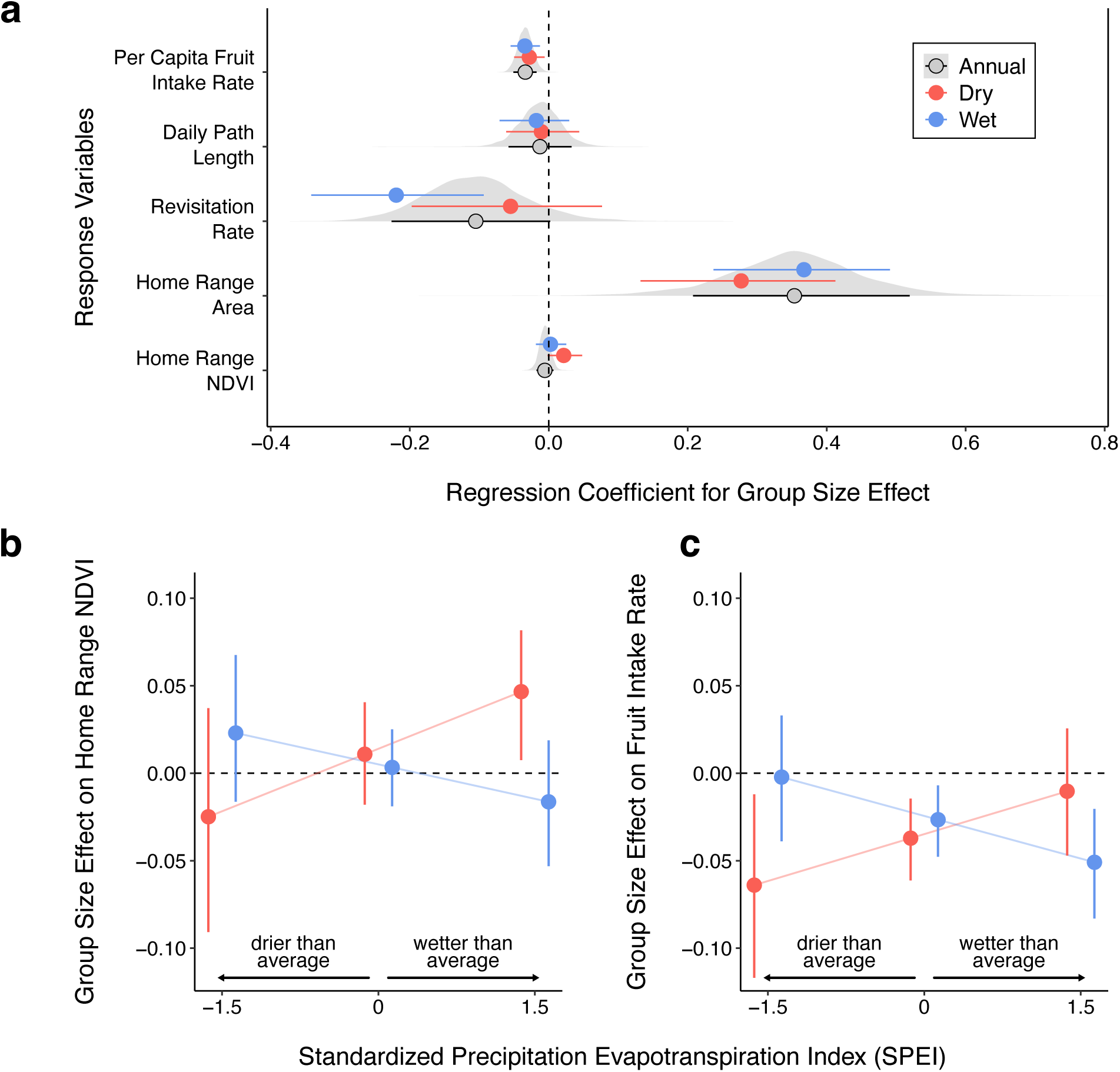
Group size effects on resource and space use. (a) Posterior estimates of the marginal effect of group size on five group-level response variables: per capita fruit intake rate, daily path length, revisitation rate, home range area, and home range normalized difference vegetation index (NDVI). Points indicate median posterior estimates, and solid lines represent 89% Highest Posterior Density Intervals (HPDIs). Blue and red point intervals correspond to wet and dry season effects, respectively, while black/grey intervals indicate annual (non-seasonal) effects. Grey density slabs show the full posterior distributions for the annual effects. (b,c) Point intervals show posterior estimates of the effect of group size on mean home-range NDVI (b) and per-capita fruit intake rate (c) conditional on representative Standardized Precipitation–Evapotranspiration Index (SPEI) values (–1.5, 0, 1.5), illustrating the *three-way interaction* between group size, season, and seasonal severity (SPEI). Points are posterior medians with 89% HPDIs (vertical bars). Blue and red denote wet- and dry-season estimates, respectively. Lines connect median effects across SPEI values to aid visual comparison. In both panels, y-axis values greater than 1 indicate a positive group-size effect (larger groups have higher NDVI/intake), and values less than 1 indicate a negative effect. On the x-axis, negative SPEI values represent abnormally dry periods and positive values represent abnormally wet periods, relative to typical conditions within each season (Jan–Apr = dry; May–Dec = wet). Fruit intake models used *n* = 335 focal individuals across 11 groups; daily path length models used *n* = 996 group-days across 11 groups; home range area and NDVI models used *n* = 156 annual and *n* = 224 seasonal home ranges across 12 groups; revisitation rate models used *n* = 101 group-years annually and *n* = 143 group-season-years seasonally across 11 groups.

### 2.2 Seasonal severity alters group-size effects on range quality and fruit intake

We quantified seasonal water-balance anomalies using the 6-month Standardized Precipitation–Evapotranspiration Index (SPEI-6). At our site, SPEI-6 is strongly correlated with the Multivariate ENSO Index (MEI; Figure S5), indicating that local hydroclimatic anomalies are largely ENSO-linked. We therefore treat SPEI-6 as our primary measure of local climate anomalies, interpreting negative values as El Niño-like (drier-than-average) and positive values as La Niña-like (wetter-than-average).

For home-range quality, larger groups generally occupied areas with higher NDVI in the dry season, but not in the wet season (Figure 2a; *β*_dry_ = 0.02[0, 0.05]; *PP >* 0 = 0.94; *β*_wet_ = 0[−0.02, 0.02]; *PP >* 0 = 0.545). This positive relationship was strongest during *exceptionally wet* dry seasons (La Niña-like; *SPEI* = 1.5) and near zero during *exceptionally dry* dry seasons (El Niño-like; *SPEI* = −1.5; Figure 2b). In contrast, during the wet season, the effect of group size on home-range NDVI was inconsistent across the SPEI gradient.

For per-capita fruit intake, group size had an overall negative effect across both seasons (Figure 2a). This negative relationship was most pronounced under climatically extreme conditions: during *very dry* dry seasons (El Niño-like; *SPEI* = −1.5) and *very wet* wet seasons (La Niña-like; *SPEI* = 1.5; Figure 2c).

When climatic anomalies counterbalanced the typical seasonal pattern— wet seasons that were unusually dry (El Niño-like; *SPEI* = −1.5) or dry seasons that were unusually wet (La Niña-like; *SPEI* = 1.5)— the effect of group size on fruit intake was near-zero.

### 2.3 Larger neighbors encroach onto smaller focal groups

To evaluate how group size asymmetries shape intergroup spatial dynamics, we modeled proportional home range overlap between neighboring group-dyads using hierarchical Social Relations Models (SRMs) [53, 54] within a causal inference framework [55]. This approach accounts for repeated measures within and across dyads by simultaneously estimating group-level tendencies (e.g., a group’s general propensity to overlap with others) and dyad-specific effects (e.g., the unique overlap between a particular pair). We incorporated group-level predictors (focal and neighbor group size) and a dyad-level predictor (interaction between focal and neighbor group size) to explain variation in asymmetric (i.e., directional) overlap (PO*_fn_*), defined as the proportional home range overlap of a focal group *f* with a specific neighbor *n*. By adjusting for the home range area of *f*, PO*_fn_* can be interpreted as the degree of encroachment by the neighboring group onto the focal group’s range. While SRMs are typically applied to dyads of individuals, extending the framework to dyads of animal groups (a) is key to understanding how within-group and between-group competition jointly shape behavior, and (b) provides a new tool for assessing group-level social networks.

Using this approach, we found that an increase in neighbor group size usually led to an increase in PO*_fn_* (Figure 3). However, this relationship was strongest when the focal group was small (∼8 monkeys; *β_P O_* = 0.61[0.25, 0.98]; *PP >* 0 = 0.995), moderate when the focal group was intermediate in size (∼21 monkeys; *β_P O_* = 0.17[0.06, 0.28]; *PP >* 0 = 0.992), and weakly negative when the focal group was also large (∼35 monkeys; *β_P O_* = −0.26[−0.65, 0.11]; *PP >* 0 = 0.128). This pattern held when examined annually and in the wet season, but not in the dry season, during which PO*_fn_* was generally reduced and neighbor group size had minimal impact (Figure 3).

**Figure 3:**
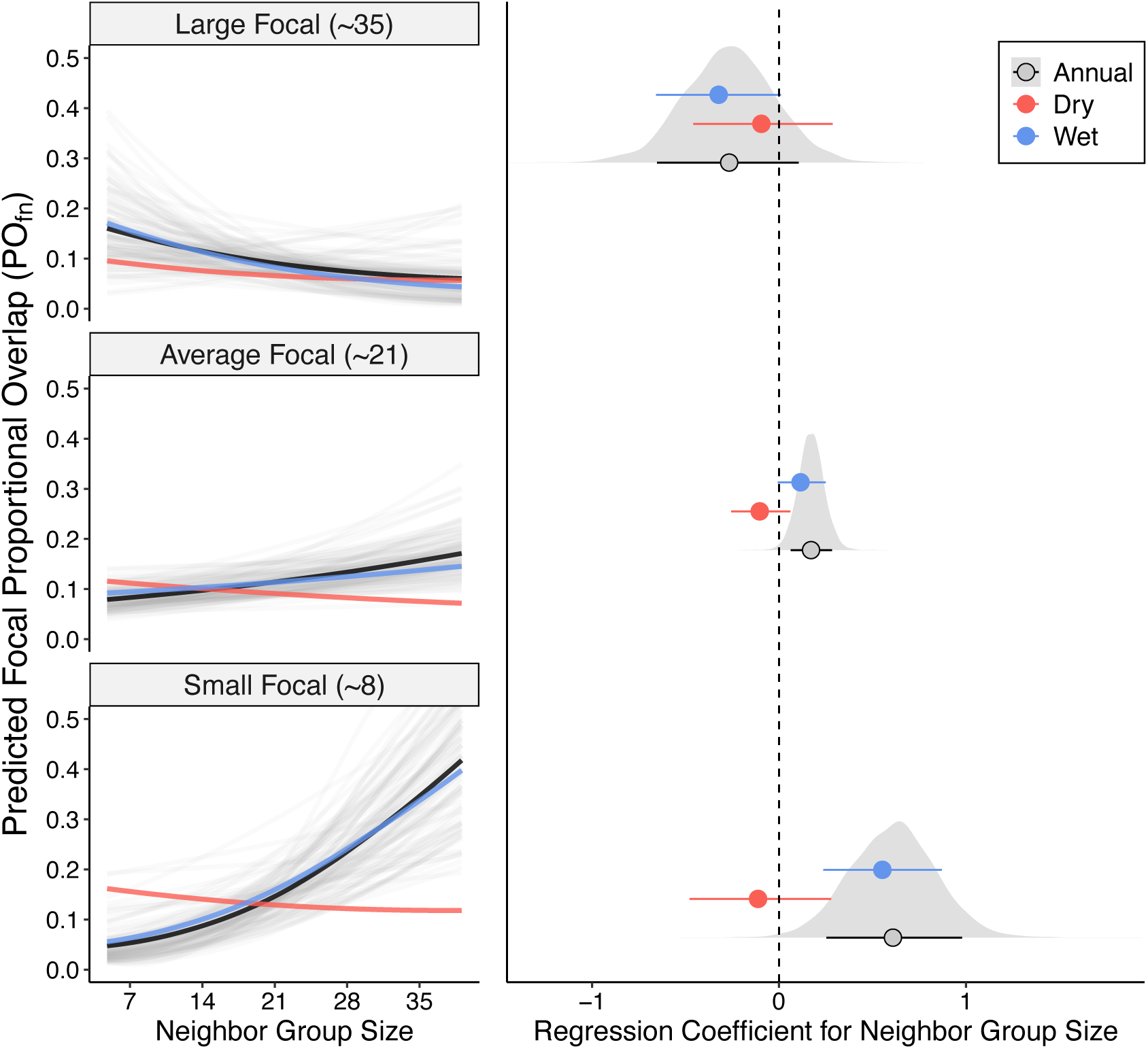
Relative group size effects on neighbor encroachment. Left panels shows posterior predictions of proportional focal home range overlap (PO*_fn_*) as a function of neighboring group size across representative focal group sizes. Light grey lines depict 100 randomly sampled posterior draws from the annual model; dark black lines show the median annual prediction; red and blue lines show median predictions for the dry and wet seasons, respectively. Right-hand plot displays median posterior effect sizes and 89% Highest Posterior Density Intervals (HPDIs) for both annual and seasonal models, corresponding to the predictions in the left panels. Grey density slabs are posterior distributions for the annual effects. The annual model was fit to *n* = 930 dyad-years (59 dyads, 12 groups); the seasonal model used *n* = 978 dyad-season-years (56 dyads, 12 groups). Each dyad contributes two observations per year or season-year to account for outcome asymmetry.

To better understand if smaller or bigger groups in a dyad drove temporal increases in overlap (i.e., change in PO*_fn_* at time *t* to *t* + *i*), we conducted a complementary analysis of dyads that showed both substantial increases in annual PO*_fn_* (from *<* 25% to *>* 45%) and shifts in relative group size (net change ≥ 5) (see section 4.4). In 84% of cases, the group that caused the overlap increase was the one that became relatively larger over time (Figure 4), consistent with a competitive advantage of larger groups for space.

**Figure 4:**
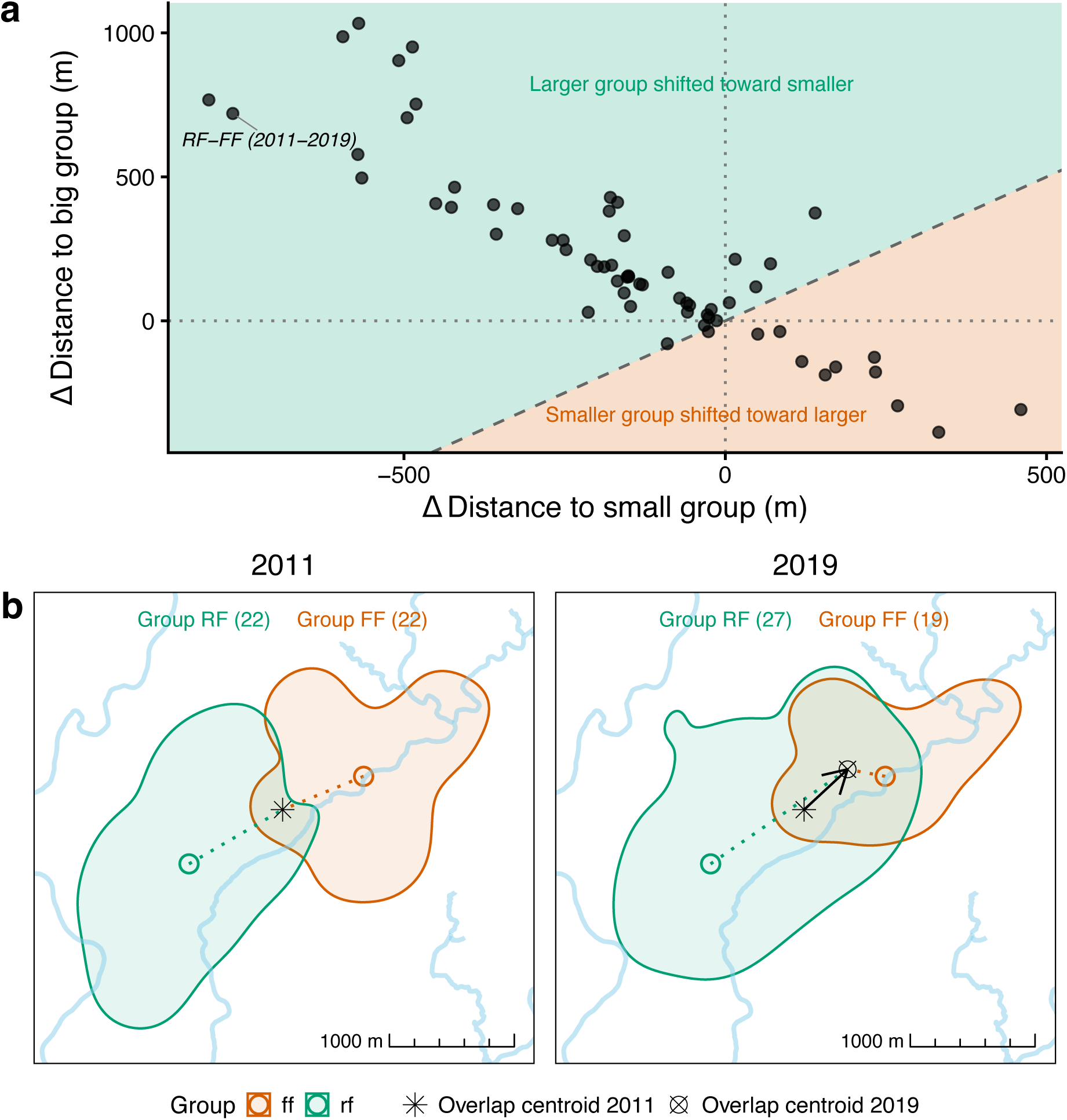
Directional shifts in home range overlap. (a) Each point represents a dyad-year pair with a marked increase in overlap and a substantial shift in relative group size. Axes show the change in distance Δ*D* from each group’s home range centroid at time *t* to the overlap centroid at time *t* + *i*, relative to the original distance at *t*. The x-axis shows Δ*D* for the group that became relatively smaller; the y-axis for the group that became larger. Positive values indicate the overlap centroid moved away; negative values indicate it moved closer. Shaded colors indicate which group drove the shift. The dashed diagonal marks equal shifts. (b) Example of dyad RF–FF (2011–2019) showing the overlap centroid’s shift. Polygons represent 95% home ranges; large circles show group centroids. Dotted lines connect group centroids to overlap centroids; the arrow shows directional change. Point shapes indicate overlap centroid locations; group sizes are in parentheses.

### 2.4 Dry conditions increase encounters and reduce overlap

To investigate how seasonality affects intergroup encounter rates, we used the ctmm framework to estimate expected space-time overlap between group home ranges (i.e., encounter rate) [56]. We then fit dyadic Bayesian multiple-membership models to assess seasonal variation in encounter rates as a function of symmetrical (i.e., non-directional) dyadic predictors: absolute group size difference, overlap zone area, and vegetation greenness within the overlap zone (measured by mean NDVI). Predicted encounter rates were consistently higher in the dry season than in the wet season, despite reduced dyadic overlap (Figure 5a). This pattern is consistent with a stronger effect of overlap zone size on encounter rate in the dry season, indicating more frequent encounters per unit of shared space compared to the wet season (posterior contrast: *β*_wet-dry_ = −0.49[−0.78, −0.16]). Encounter probability was highest in overlap zones with greater vegetation productivity, particularly in the dry season (*β*_HRQ_ = 0.42[0.09, 0.74]). In contrast, absolute group size difference had no consistent effect on encounter rate across both scales (Figure 5c).

**Figure 5:**
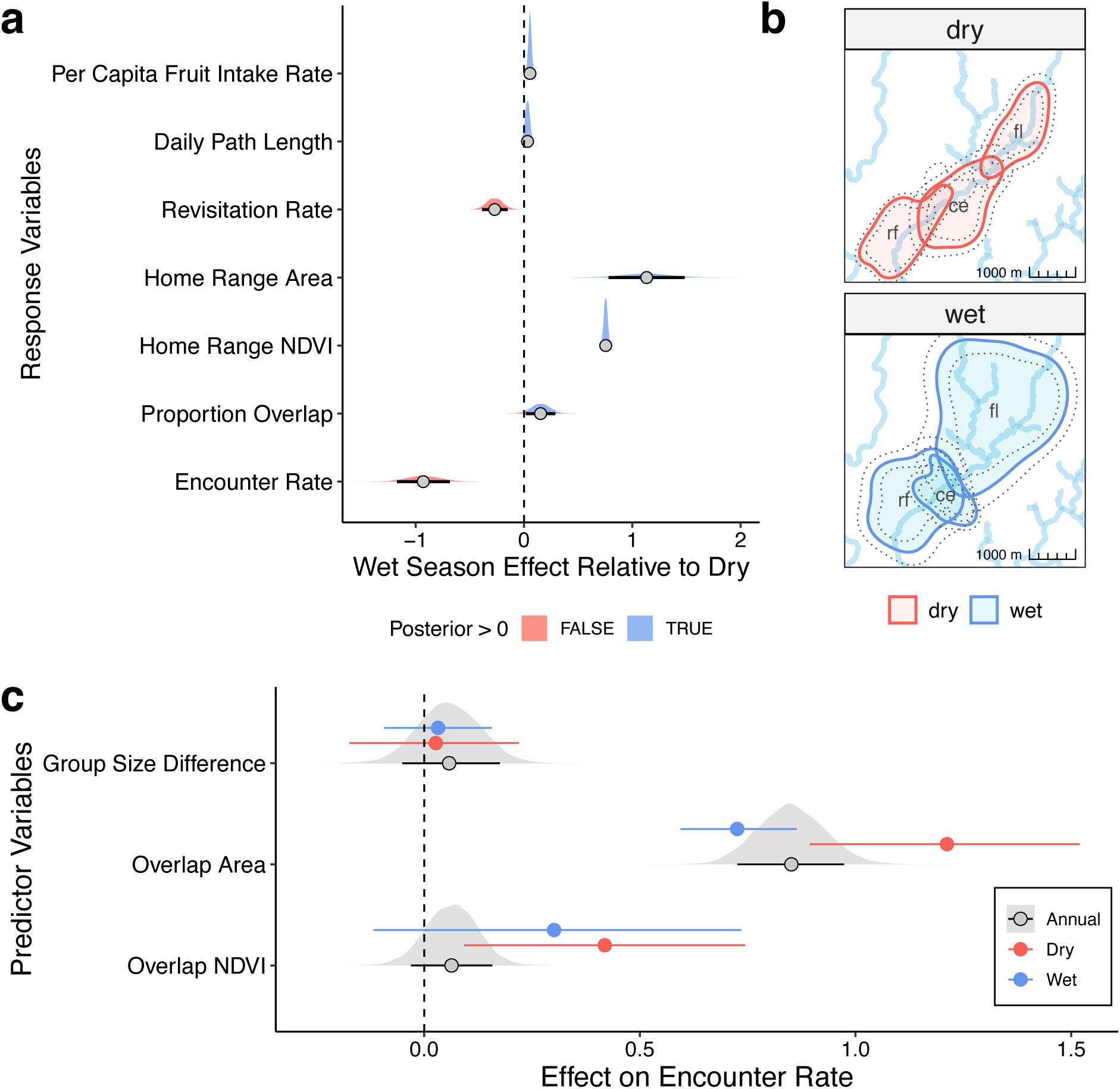
Seasonal changes in fruit intake, ranging behavior, and encounter rates. (a) Posterior densities, medians, and 89% Highest Posterior Density Intervals (HPDIs) for wet vs. dry season contrasts across all seven response variables: fruit intake, path length, revisitation rate, home range size, home range NDVI, proportional overlap (PO*_fn_*), and encounter rate. Red shading indicates higher values in the dry season; blue shading indicates higher values in the wet season. (b) Seasonal comparison of home range overlap among three groups in 2018. (c) Posterior medians and 89% HPDIs for annual and seasonal effects of group size difference, overlap size, and overlap NDVI on encounter rate. Blue and red point intervals correspond to wet and dry season effects, respectively, while black/grey intervals indicate annual (non-seasonal) effects. Grey density slabs are posterior distributions for the annual effects. Sample sizes for fruit intake, path length, revisitation rate, and home range models are as reported in Figure 2; overlap model sample sizes are as reported in Figure 3. Encounter rate models were fit to *n* = 269 dyad-years (42 dyads, 11 groups) annually and *n* = 280 dyad-season-years (41 dyads, 11 groups) seasonally. Note that seasonal and annual models are fit to different subsets of the data, which may contribute to discrepancies between seasonal and annual effect sizes.

## 3 Discussion

Our findings show that trade-offs of group living are shaped by complex interactions between environmental conditions and group size, which jointy mediate how within- and between-group competition influence the social and spatial structure of a population. Larger capuchin groups showed reduced per-capita fruit intake efficiency, indicating stronger within-group competition over clumped resources— consistent with the Ecological Constraints Model [8]. However, this increased competition did not translate into higher daily travel costs for larger groups (contra common expectation; e.g., [11, 12]). Instead, they appear to mitigate foraging competition by leveraging their numerical advantage to outcompete smaller groups, gaining access to *less frequently visited* areas during the wet season (when food is abundant and dispersed) and to *higher-quality* patches during the dry season (when resources are concentrated in riparian zones).

The transition from wet to dry season may mark a shift from indirect to more direct between-group competition, as resources become limited and more economically defensible. Supporting this, we found that home range overlap declines in the dry season— consistent with increased active defense [57]—while intergroup encounter rates rise, particularly in areas with higher vegetation productivity near permanent water sources. This pattern matches theoretical models predicting that frequent encounters can drive spatial separation, as repeated costly interactions discourage intrusion into contested areas, thereby promoting more fixed home-range boundaries and reducing resource loss to neighbors [58, 59]. Reduced overlap and the tendency for larger groups to occupy more productive ranges in the dry season suggest that increased encounters may enable them to secure more exclusive access to the highest-quality areas through the exclusion of smaller groups. While direct evidence is lacking for whether larger groups gain a net advantage from the wet-dry season transition, our results underscore how short-term interactions can have long-lasting ecological consequences on space use and access to resources.

These findings parallel predictions from the resource dispersion hypothesis (RDH), originally developed to explain group formation in territorial social carnivores [39, 60]. In these species, individuals that might otherwise occupy separate territories form groups when resources are sufficiently rich and dispersed to support multiple individuals within a single territory [17, 61]. This can promote sociality even in species with little cooperative behavior or predator-defense needs [34]. Although social primates like capuchins differ in many respects— including more rigid social structures [62], additional drivers of sociality (e.g., alloparental care [44], social learning [63]), and generalist diets [64]— our findings suggest that similar ecological dynamics may still emerge. When the environment can support more individuals or groups, capuchins appear to reduce investment in active defense against neighbor intrusion.

The RDH alone cannot explain capuchin sociality as groups remain hostile during intergroup encounters regardless of resource dispersion [51]. Yet, it may help explain how smaller groups persist on the landscape despite overlapping with multiple more powerful neighbors. For instance, our findings indicate that larger groups tend to avoid overlapping with other large groups, perhaps to reduce the risk and energetic cost of conflict with similarly powerful rivals, where outcomes are less predictable [65–67]. This pattern aligns with game-theoretic models of animal contests, which predict that when opponents are evenly matched, mutual assessment and conflict avoidance strategies evolve to reduce costly escalations [68, 69]. Such mutual avoidance may inadvertently benefit smaller groups by creating “buffer zones”— underused areas that smaller groups can exploit while remaining undetected [70]. Comparable patterns are seen with predator-prey dynamics: white-tailed deer (*Odocoileus virginianus*), for example, are more likely to occur in areas where wolf (*Canis lupus*) pack territories overlap [71, 72]. Similarly, during large-scale human conflicts, activity often declines near war zone buffer areas, leading to natural reforestation and surges in wildlife populations and biodiversity [73, 74]. Our results suggest that when resources are sufficiently abundant and dispersed to support multiple groups within a given area, spatial overlap increases— especially between groups with strongly asymmetric competitive abilities. Such conditions allow dominant groups to access new resources with minimal conflict while enabling smaller groups to persist in underused zones between the ranges of their more powerful neighbors.

Our results show that greater within-group competition with increasing group size does not necessarily lead to higher daily travel costs, even for species that primarily forage on clumped resources. Folivores are known to avoid this trade-off due to the relatively widespread and homogeneous nature of their food resources (the so-called folivore paradox) [40, 75]. Dietary generalists may achieve the same effect by shifting to more readily available resources like insects [76]. Beyond such fallback strategies, our findings suggest that larger groups can also alleviate foraging pressure by expanding their range, increasing the diversity of fruiting trees exploited, and lowering revisitation frequency. This strategy likely enables access to less-depleted, higher-quality patches, particularly along range peripheries where fruiting trees contain more ripe fruit [77]. Considering physiological constraints on daily energy expenditure [78, 79], rotating among foraging areas over longer timescales may be a more efficient strategy than increasing daily travel distance (see subsection S1.7). Further, our findings show that climatic extremes (i.e., exceptionally severe dry or wet seasons) exacerbate within-group competition for large groups, leading to decreased per-capita foraging efficiency. This pattern suggests that extreme climates compound ecological constraints on larger groups. Tropical forests are particularly sensitive to ENSO-driven anomalies [47, 80], which have well-documented fitness consequences for mammals (including capuchins). El Niño events that prolong or intensify the dry season increase heat stress and reduce survival and reproductive success [49, 81], whereas La Niña events that amplify wet-season rainfall disrupt phenological and arthropod cycles, occasionally triggering famine and widespread mortality [82, 83]. Such extremes likely impose additional constraints on large groups, especially if they suppress overall food production so that even the best-quality patches cannot support many consumers.

By contrast, intermediate climatic anomalies (e.g., ENSO-linked conditions that counterbalance the typical seasonal cycle) appear to relax these constraints and may even enhance between-group advantages for large groups. During wetter-than-average dry seasons and drier-than-average wet seasons, group size had little to no effect on per-capita foraging efficiency, and larger groups tended to occupy higher-quality ranges. One possibility is that such conditions increase spatial heterogeneity in habitat quality (analogous to the Intermediate Disturbance Hypothesis [84]), creating patchier resource landscapes that larger groups can more effectively monopolize— a pattern consistent with broader evidence that climate variability and resource heterogeneity selects for enhanced cooperation and resource defense strategies [85–87]. Thus, in highly heterogeneous environments, between-group competition may have played a greater role in the evolution of sociality than often appreciated [7, 88]. Future work that explicitly quantifies the distribution of key resources (beyond vegetation greenness) and links group size to fitness would clarify the ecological conditions that favor within-versus between-group competition. Additional insight into the evolution of sociality may come from integrating these trade-offs within a multilevel selection framework [89, 90].

We demonstrate that the benefits of between-group dominance and the costs of within-group competition vary predictably across changing environments. Such shifting trade-offs may help explain why group sizes vary widely within populations and over time. Small groups can persist by (a) having reduced within-group foraging pressure, (b) rooting themselves spatially to avoid conflict with neighbors, and (c) maintaining a competitive advantage within their core area regardless of opposing group size [37]. In contrast, large groups can endure by asserting spatial dominance and monopolizing the most valuable resources which could act as a buffer against intensified within-group competition in the face of resource unpredictability. Nevertheless, this buffering capacity may have limits under particular social or ecological conditions (e.g., prolonged climatic extremes), potentially explaining the rare occasions when large groups permanently fission [91]. Unlike ‘facultative groupers’ (i.e., fission–fusion species) that can flexibly adjust group size to local circumstances [31], ‘obligate groupers’ such as capuchins are bound by the long-term benefits of group living (e.g., cooperative breeding, protection from predators or infanticidal conspecifics [92]), which regularly outweigh the costs (e.g., within-group competition, disease transmission) [29]. For such species, permanent fission represents a last resort when groups approach the upper limit of viable size under prevailing conditions [93], with the potential to fundamentally alter the balance of power across the social landscape.

Understanding how group size influences resource competition is inherently complex. Many studies analyze within- and between-group dynamics in isolation, overlooking how they interact and vary with ecological context. Our study addresses this gap by integrating multi-group longitudinal data with a novel, spatially explicit extension of the social relations model. This framework quantifies how both focal and neighboring group traits shape ranging behavior, revealing whether competitive pressures are specific to certain group pairs or reflect broader, population-level patterns. A remaining obstacle, however, is accounting for unmonitored groups— a common limitation in long-term field studies [23]. Adopting a more holistic “neighborhood” perspective— akin to the distinction between local and global dispersal [94, 95]— could offer deeper insight by integrating the combined effects of multiple neighbors and inferring impact of unobserved groups. Our framework provides the foundation for such extensions, and for a more nuanced consideration of underlying social dynamics (e.g., age–sex structure, dominance hierarchies, collective action problems [96]).

Our study reveals that climate and demographic change shape the trade-offs linking within- and between-group competition. While within-group competition imposes an upper limit on group size [9], and between-group competition and predation likely set a lower limit [10], the strength of these constraints is context-dependent. Their relative influence shifts with climate and resource availability [38], allowing groups of different sizes to coexist, each benefiting from distinct ecological and social conditions. Yet, increasingly intense and erratic ENSO cycles, together with ongoing habitat fragmentation, are rapidly altering tropical landscapes [80, 97–99], making resource availability increasingly unpredictable. Such climatic fluctuations appear to amplify within-group competition under some ecological contexts (e.g., El Niño dry seasons) while enhancing between-group advantages under others (e.g., La Niña dry seasons). These findings raise the question of whether ongoing climate change will destabilize these trade-offs, ultimately tilting the balance in favor of smaller or larger groups in the future.

## 4 Methods

### 4.1 Data Collection

The Lomas Barbudal Monkey Project dataset contains longitudinal records from 13 habituated groups of white-faced capuchins, spanning 1990 to 2025. These records include data on demographics, foraging, social interactions, and group movement. The 13 study groups have varying observation periods, reflecting differences in when each group was introduced into the study or formed through permanent fission from an existing study group (Figure 1c). When a permanent fission occurs, the group that retains the natal home range (invariably the larger faction) keeps the original identifier of the parent group, whereas the group that splits off and forms a new home range is given a new identifier. Here, we draw on data from 12 of the 13 study groups, as one group (DT) was excluded due to insufficient location data.

Location data from 1991 to September 2009 (none available for 1990) consist of georeferenced sleep-site records digitized from field notes originally used to relocate groups each morning. From September 2009 to 2025, handheld GPS units (Garmin GPSmap 62s, 64, 64s, 66sr) were clipped to or placed in researchers’ backpacks and used to record both sleep-site locations and daily movement trajectories as researchers followed the monkeys. GPS tracking data have been cleaned and processed through 2020, and GPS-derived sleep-site locations through 2023. Sleep-site data therefore span from the start of 1991 to the end of 2023, whereas GPS tracking data extend from September 2009 to April 2020 (Figure S1). Previous work has validated the reliability of both data types for estimating home ranges [43, 100].

Researchers positioned themselves near the group’s center when not conducting focal follows. GPS units recorded locations at intervals of five minutes (2009-2012) or 30 seconds (2013-2020). We calculated median annual and seasonal group sizes from 1991 to 2023 using daily group censuses with at least six hours of observation to maximize the likelihood of detecting all individuals present. We summarized fruit foraging behavior from focal follows conducted from July 1, 2006 to June 30, 2021. These records included the number of bites of fruit observed to be ingested, the duration each individual was visible, and the age and sex of the focal individual. Further specifics on data collection and processing are provided in Jacobson *et al.* [100] and Perry *et al.* [101].

### 4.2 Environmental Data

The tropical dry forests in and around the Lomas Barbudal Biological Reserve experience strong annual and seasonal variation in vegetation phenology, affecting the availability of key resources such as food, water, and shade for capuchins. To quantify interannual and interseasonal variation in vegetation phenology across the study period, we extracted annual and seasonal Normalized Difference Vegetation Index (NDVI) values from 1991 to 2023 using satellite imagery extracted from Google Earth Engine [102]. Although NDVI can be an imperfect proxy for food availability in tropical forests [103], it has been shown to measure long-term habitat variability well in this region [104], particularly in the dry season when resources such as water, fruit, and shade are concentrated in greener riparian zones. NDVI quantifies the “greenness” of each pixel in a given image, serving as a proxy for vegetation health and abundance. We defined a bounding box around the Lomas Barbudal Biological Reserve and surrounding forest as the spatial extent for all image processing. We used surface reflectance data from Landsat 5, Landsat 7, and Landsat 8 satellites to obtain red (RED) and near-infrared (NIR) bands required for NDVI calculation. To remove cloud-contaminated pixels, we applied a mask using the quality assessment (QA_PIXEL) band. NDVI was calculated for each image as:

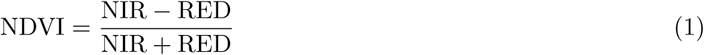

For each year and season, we created pixel-wise maximum composites to reduce noise from residual cloud contamination and obtain the best quality pixel value. Seasonal windows were defined as February–April (peak dry season) and September–November (peak wet season). These seasonal and annual median NDVI images were clipped to the study area and exported at 30-meter resolution (see subsection S1.3 for details). In addition to NDVI, we extracted rasters of tree cover from the Hansen Global Forest Change dataset, [105], which were later used to refine home range estimates (see section 4.3). All processing of remote sensing data was conducted using Google Earth Engine’s Python API, and rasters were exported as .tif files for further analysis in R [106].

We quantified seasonal severity using the Standardized Precipitation–Evapotranspiration Index (SPEI), which measures anomalies in water balance calculated as precipitation minus potential evapotranspiration (PET)—the amount of water lost to evaporation and plant transpiration under existing temperature and radiation conditions. Monthly SPEI values (1990–2023) were obtained from the ERA5-Drought reanalysis dataset [107] for the Lomas Barbudal Biological Reserve and surrounding areas used by our study population. SPEI values were standardized within seasons, such that negative values indicate abnormally dry periods and positive values indicate abnormally wet periods relative to the typical range within that season (Jan–Apr = dry; May–Dec = wet). We used a 6-month accumulation period, which captures broader climatic conditions and aligns most closely with ENSO variability (Figure S5; see subsubsection S1.3.2 for comparison with 1-month SPEI).

### 4.3 Group-level Response Variables

For several of our questions of interest, our observation and response variables of interest were properties of individual groups (Table 1).

#### Quantifying fruit intake and daily path length

Per capita fruit intake rates were estimated from focal follow data collected between July 1, 2006, and June 30, 2021. This dataset includes 4,952 hours of usable observation time from 335 individually identified focal individuals. For each focal individual per day, we recorded the number of bites of fruit ingested, rather than whole fruits, as many fruits require multiple bites to consume. Intake data were restricted to periods when both the mouth and hands of the individual were visible, ensuring accurate detection of fine-scale foraging behavior.

To quantify daily energy expenditure, we used GPS-tracking data to generate 996 daily path lengths across 11 groups from September 2009 to April 2020, applying the continuous-time speed and distance (CTSD) method described by [108] using the ctmm R package [109]. This approach estimates daily path length by multiplying the estimated mean speed from a fitted daily continuous-time movement model by the total daily sampling duration. By separating the continuous-time movement process from the discrete-time sampling process, CTSD is less sensitive to sampling rate, path complexity, and location error than traditional methods based on totaling step lengths. We included only GPS tracks with at least 11 continuous hours of data per day, corresponding to minimum daylight hours, as capuchins rarely move at night.

#### Quantifying home range area, greenness and revisitation rate

We used sleep-site data from 1991 to 2023 to estimate home range area (validated in [43]), enabling us to use the full 33-year dataset, including years pre-dating GPS, and capture the widest range of demographic changes and study groups over time. Our validation study demonstrated that 98% utilization distributions (UD) from sleep ranges accurately aligned with the conventional 95% UD from GPS-tracking data as capuchins typically refrain from sleeping in the far outskirts of their home range [43]. Based on this, we used 98% UDs to estimate home range area, representing the smallest area with a 95% probability of finding the group considering its total movements.

We calculated 156 annual and 224 seasonal sleep home ranges across 12 groups using auto-correlated kernel density estimation (AKDE) implemented using the ctmm R package [109]. AKDE estimates the UD by fitting a series of continuous-time movement models to the location data, selecting the best-fitting model based on the data’s autocorrelation structure, which informs the kernel smoothing bandwidth and yields home range estimates that account for location error and irregular sampling [110, 111]. Seasonal ranges were segmented as January–April (dry) and May–December (wet), based on behavioral shifts in daily distance to rivers, rather than on climatic seasonality, which typically includes December in the dry season (Figure 1). Only data segments suitable for home range estimation, identified via visual inspection of variogram regression plots, were included in downstream analyses [112]. We also fit these home ranges using an integrated resource selection function (iRSF) of tree cover following [113]. This imposed “soft barriers” by down-weighting areas without trees, like pasture-lands, which home range estimates may expand into but capuchins rarely use. We measured home range greenness by extracting the mean NDVI value within each group’s mean home range boundary for the corresponding year or season using the terra package [114].

We quantified revisitation rates using the revisitation() function in the ctmm package, which provides a continuous-time, model-based estimate of how frequently a capuchin group returns to previously used areas. After fitting a continuous-velocity movement model and estimating each group’s AKDE utilization distribution (UD), revisitation() constructs a revisitation-rate surface on the same spatial grid as the UD. At each grid cell, the local revisitation rate is determined by the product of the model-estimated instantaneous speed and the UD value at that location. This represents the expected frequency with which the animal’s path re-enters a small circular neighborhood around that point, scaled by the grid’s radial resolution. Averaging this revisitation-rate surface over the UD yields a single mean revisitation rate per radial distance for each group, which describes the typical frequency with which a group revisits locations within its home range (that is, how often the group returns to previously used areas versus moving into less-used areas). The revisitation rate is defined per unit meter by default, but can be multiplied by any larger distance (e.g., 25–50 m) to obtain an expected revisitation rate for that spatial scale. Because ctmm derives revisitation from a continuous-time model rather than from discrete visit/leave thresholds, the estimate is insensitive to the choice of revisitation radius and robust to the sampling schedule of the location data (provided the data resolve the movement’s velocity timescale). We used the GPS-tracking dataset (September 2009 - April 2020) to estimate revisitation rates, which were calculated separately for each group-year and group-season, matching the observational scale used for UD estimation. Although narrower in longitudinal scope than our sleep-site records (1990 - 2023), these tracking data provide the spatial and temporal resolution needed to estimate velocity, generate accurate UDs [43], and capture diurnal space-use patterns relevant to revisitation. An empirical comparison with trajectory-based revisitation estimates from the recurse package [115] is provided in subsection S1.8.

### 4.4 Dyad-level Response Variables

For other questions of interest, our observation and response variables of interest were properties of dyads of groups— a unique pairing of groups (Table 1). This distinction between group- and group-dyad-level outcomes is important for accurate statistical estimation.

#### Quantifying proportional overlap and range shifts

We measured home range overlap between group-dyads among known habituated groups throughout the study period. We calculated the proportion of a group’s home range that overlaps with a specific neighbor (PO*_fn_*). This approach allows us to assess the degree to which a neighbor can encroach into a focal group’s range, and the potential loss of a group’s available resources to that neighbor. PO*_fn_*, was calculated by:

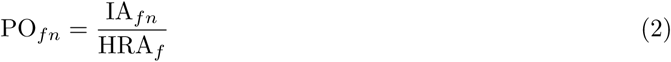

Here, IA*_fn_* represents the area of intersection between the focal and neighbor group’s home ranges, and HRA*_f_* is the home range area of the focal group. PO*_fn_* measures an asymmetric group-dyad outcome. To demonstrate, imagine that group A and group B have home ranges with some intersection, IA*_ab_*. While IA*_ab_* = IA*_ba_*, PO*_ab_ ≠* PO*_ba_* because HRA*_a_ ≠* HRA*_b_*. Using this asymmetric variable, we focus on the proportional overlap from the focal group’s perspective. For every observation, the neighboring group also has a PO*_fn_* measurement but the groups are switched.

While PO*_fn_* captures asymmetric space use and can suggest which group is being encroached upon at a given time, it does not track the directionality of range shifts *over time*. To assess whether larger or smaller groups drove increases in overlap, we conducted a complementary analysis on dyads that exhibited both a substantial increase in spatial overlap (from *<*25% to *>*45%) and a clear change in relative group size (absolute difference ≥5 individuals). These thresholds were chosen to isolate cases with meaningful shifts in both space use and relative group size, avoiding ambiguity in assigning which group is considered the “big” group (e.g., a group may remain larger overall but shrink more than its neighbor from time *t* to time *t* + *i*) and minimizing noise from minor fluctuations. For each of these dyads (63 cross-year comparisons across 11 unique group pairs), we measured the distance from each group’s home range centroid at time *t* to the centroid of the overlap zone at both time *t* and time *t* + *i*, and calculated the change in distance (Δ*D*):

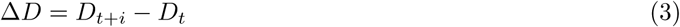

A positive Δ*D* indicates that the group shifted toward the other group in the dyad, while a negative value indicates the other group shifted toward them. The group with the larger magnitude of Δ*D* was considered the one that drove the increase in overlap.

#### Quantifying encounter rates

We computed encounter rates using the ctmm::encounter() function, following [56], to quantify each dyad’s expected space–time overlap. As with revisitation rates, we used UDs derived from GPS-tracking data (2009–2020) to better capture fine-scale, diurnal space use. Importantly, ctmm::encounter() does not count empirical instances in which groups simultaneously enter a defined encounter radius. Rather, it estimates how often two groups’ modeled space use would place them within a small spatial radius at the same moment, given their long-term movement patterns and assuming stationary movement parameters (e.g., range crossing time and autocorrelation structure). Using this framework, we define the encounter rate ER*_ij_* as the relative probability that, at any given moment, groups *i* and *j* are found within the same 1 m^2^ area (a scale that can be adjusted to any encounter radius). These values are relative and interpretable by comparing across dyads; for example, if ER*_ab_* = 0.4 and ER*_ac_* = 0.2, then groups *a* and *b* are twice as likely to encounter each other as groups *a* and *c*. Unlike PO*_fn_*, ER*_ij_* is symmetric, yielding the same value regardless of which group is labeled *i* or *j*, as group identity plays no directional role in this metric. We also validated that these model-based encounter rates correspond closely with observed inter-group encounters (see subsection S1.8).

### 4.5 Statistical Analyses

#### 4.5.1 Group-level models

We used a suite of GLMMs to examine the relationship between group size and several within-group outcomes: per capita fruit intake rate, daily path length, revisitation rate, home range area, mean home range NDVI. All models were fit in a Bayesian framework using the brms package [116]. We specified Gamma likelihoods for all models except the home range NDVI models, which used a Beta distribution, and the fruit-intake models, which used a negative binomial distribution. Group size was standardized, and additional covariates were included where appropriate based on causal assumptions outlined in our directed acyclic graphs (subsection S1.4). All models included varying effects by group, with additional varying effects by individual (for fruit intake) or by year (for home range NDVI), where appropriate. The fruit-intake models also included an offset term accounting for observation time (seconds in view). Models for home range area and daily path length included error terms on the response to account for measurement uncertainty. Because key resources regenerate annually [117] but also fluctuate between wet and dry seasons, each model was run in two versions: one including seasonality (via an interaction between group size and season), and one without. While only home range area and NDVI were explicitly measured at seasonal or annual scales, we refer to the non-seasonal models for all outcomes as “annual” for consistency. To assess how climatic anomalies modulate these dynamics, the seasonal models for home-range NDVI and fruit intake rate (proxies for between- and within-group competition, respectively) were extended to include SPEI as a three-way interaction with group size and season. SPEI was standardized within seasons, allowing us to estimate how deviations from typical wet- or dry-season conditions modify the effects of group size without introducing collinearity between SPEI and season.

#### 4.5.2 Dyad-level models

To examine how relative group size shapes between-group interactions, we used an extension of the hierarchical Social Relations Model (SRM) [53]. SRMs separate group-level and dyadic-level variation while modeling their correlation, enabling efficient partial pooling and improving parameter estimation [54]. While traditionally applied to social behavior among individuals or human households [e.g., 118–120], we extend this framework to spatial interactions between animal social groups.

We modeled proportional overlap (*PO_fn_*) using a zero-augmented (hurdle) SRM implemented in the rethinking package [121]. The hurdle component separates the probability of non-zero overlap (Bernoulli distribution) from the degree of overlap when present (Beta distribution). Key predictors included focal and neighbor group size (as group-level fixed effects) and their interaction (as a dyad-level fixed effect). Seasonal models included a three-way interaction between season, focal group size, and neighbor group size. To account for non-independence across dyads, we included varying effects for focal group ID, neighbor group ID, and dyad ID, modeled jointly via a multivariate distribution with a shared correlation structure. This structure allows, for instance, inference about whether a group that overlaps more with one neighbor also overlaps more with others— capturing cross-dyad patterns more flexibly than standard random effects. See subsection S1.5 for full statistical notation of our SRMs.

We modeled encounter probabilities using a related set of Gaussian-distributed dyadic models (after log-transforming the response), implemented in brms. While based on the same dyadic framework, these models used a different varying effects structure than the SRM described above (sensu [118, 122]), omitting correlated focal–neighbor effects. This was appropriate because encounter rate is a symmetric response—interactions between groups *i* and *j* are equivalent regardless of order— making the focal–neighbor distinction neither identifiable nor meaningful. To reflect the symmetrical nature of the data, we included two sources of random variation: a dyad-level effect, capturing dyad-specific tendencies in encounter rates, and a group-level multiple membership structure, allowing each observation to be jointly attributed to both groups involved. This approach preserves the exchangeability of group identities within dyads and appropriately partitions variance between individual group tendencies and unique dyadic relationships. All predictors were also symmetric: group size difference, overlap zone area, and vegetation greenness in the overlap zone (measured by mean NDVI). As with the home range and path length models, we accounted for uncertainty in encounter rates using measurement error modeling. To assess seasonal variation in encounters, we included separate interaction terms between season and each predictor— testing whether the influence of group size asymmetries and habitat characteristics on intergroup encounters varied by season.

We applied weakly regularizing priors to all model parameters. All models showed good convergence (*R̂* ≈ 1.00), adequate effective sample sizes, and reliable fit based on posterior predictive checks. We summarize parameter estimates using 89% Highest Posterior Density Intervals (HPDIs) and report the proportion of the posterior greater than zero (PP*>*0), which represents the posterior probability that an effect is positive (values near 0 indicate strong support for a negative effect). Following McElreath [121], we use 89% intervals not as significance thresholds but as a convention to represent uncertainty while discouraging dichotomous interpretation of results (i.e., “significant” vs. “non-significant”). See subsection S1.4 for additional model details and assumptions using a causal inference framework.

## 5 Data availability

Location data are restricted to protect the precise whereabouts of habituated, threatened primates vulnerable to the pet trade. They are archived in a limited-access Movebank repository, and available upon reasonable request to SEP. All demographic and behavioral data are openly available on Dryad [123].

## 6 Code availability

All code necessary to reproduce our analyses are openly available on Edmond [124].

## 7 Ethics statement

The study was purely observational, with GPS units carried by observers rather than attached to animals. Research protocols were approved by UCLA’s Animal Care Committee (protocol 2016–022), and all required permits from SINAC and MINAE (the Costa Rican agencies overseeing wildlife research) were secured and renewed every six months. The most recent authorizations include scientific passport #012-2024-ACAT and permit Resolución #M-P-SINAC-PNI-ACAT-0010-2024. All procedures complied with the Animal Behavior Society’s Guidelines for the Use of Animals in Research [125].

## Supporting information

Full Supplementary Information

## 8.#Acknowledgments

Thanks to Kate Tiedeman and Alison Ashbury for feedback on remote-sensing and writing respectively. Thank you to Cody Ross and Jeremy Koster for feedback on model structure, and to Richard Berl for establishing the GPS protocols used for location data collection at Lomas (see subsection S1.1 for list of all contributing field assistants). Thanks to Don Cohen for assistance with maintaining the LBMP database. We thank the Costa Rican Park Service (SINAC, Área de Conservación Arenal Tempisque) for permission to work in Lomas Barbudal Biological Reserve, and the private landowners who have granted us permission to work on their land (especially Hacienda Pelon, Brin d’Amor and the community of San Ramon de Bagaces). We thank three anonymous reviewers and the editor for their valuable feedback.

Data collection was funded primarily by grants to SEP from the National Science Foundation (grants BCS-1919649, BCS-1638428, BCS-0613226, BCS-848360, DDIG 1232371 (co-PI I Godoy), 9633991, SES9870429 and an NSF graduate fellowship), the National Geographic Society (grants 7968-06, 8671–09, 9058-12, 9795–15, 45176R-18 and a grant with co-PI B Smuts), the L.S.B. Leakey Foundation (9 grants), the Templeton World Charity Foundation, Inc. (grant 0208), 2 Wenner-Gren grants (one with co-PI I. Godoy), various UCLA COR grants and internal grants from the University of Michigan, Sigma Xi, 5 years of funding by MPI-EVAN, and logistic support from the Wild Capuchin Foundation. BJB supplemented data collection via funds from the American Society of Primatologists, ARCS Foundation and an NSF GRF (grant no. 1650042). This work was supported by the Alexander von Humboldt-Stiftung through funds awarded to MCC.

## 9 Author Contributions

We use the Contributor Roles Taxonomy (CRediT) to detail author contributions: Conceptualization (OTJ, BJB, MCC, SEP); Data curation (OTJ, SEP, BJB); Formal Analysis (OTJ, BJB); Funding acquisition (SEP, MCC, BJB); Investigation (SEP, OTJ, BJB); Methodology (OTJ, BJB, SEP, GEF); Project administration (SEP, BJB, OTJ); Resources (SEP, BJB, MCC); Software (OTJ, BJB, GEF); Supervision (BJB, SEP, MCC, GEF); Visualization (OTJ); Writing – original draft (OTJ, BJB, SEP); Writing – review & editing (OTJ, SEP, BJB, GEF, MCC).

