## Supplementary material for "Environmental fluctuations alter the competitive trade-offs of group size in a social primate": Full Supplementary Information

### S1.1 Acknowledgements of field assistants:

The following people assisted SEP, BJB and OTJ in data collection: J. Anderson, C. Angyal, L. Appleby, K. Atkins, A. Autor C., M. Bergstrom, R. Berl, L. Beaudrot, T. Bishop, A. Bjorkmann, L. Blankenship, T. Borcuch, J. Broesch, A. Büry, D. Bush, J. Butler, F. Campos, C. Carlson, S. Carnegie, S. Caro, L. Chuaqui, A. Cobden, C. Collins, G. Corradini, M. Corrales, J. Damm, B.A. Davis, C. deRango, C. Dillis, N. Donati, G. Dower, R. Dower, A. Duchesneau, K. Feilen, J. Fenton, S. Fiello, K. Fisher, A. Fuentes J., M. Fuentes, T. Fuentes A., A. Gaston, C. Gault, H. Gilkenson, M. Glenwright, I. Godoy, I. Gottlieb, J. Griciute, J. Gros-Louis, L.M. Guevara R., M. Guimond, L. Hack, M. Hammel, R. Hammond, R. Hamrick, A. Hanadari-Levy, S. Herbert, C. Hirsch, M. Hoffman, A. Hofner, C. Holman, J. Hubbard, S. Hyde, M. Jackson, S. Jackson, E. Johnson, K. Kajokaite, M. Kay, E. Kennedy, D. Kerhoas-Essens, S. Kessler, D. Khieninson, P. Kolence, W. Krimmel, W. Lammers, M. Lechner, S. Lee, S. Leinwand, L. Johnson, S. Lopez Plaza, T. Lord, S. MacCarter, J. Mackenzie, F. McKibben, J. Manson, M. Mayer, W. Meno, A. Mensing, M. Milstein, A. Mitchell, C. Mitchell, W. Meno, J. Mudde, Y. Namba, D. Negru, A. Neyer, C. O'Connell, J.C. Ordoñez J., F. Ouweleen, N. Parker, B. Pav, S. Pereira, K. Perry, J. Pinnock, R. Popa, K. Potter, K. Ratliff, K. Reinhardt, N. Roberts B., E. Rothwell, J. Rottman, H. Ruffler, S. Sanford, C.M. Saul, S. Schading, I. Schamberg, S. Schembari, N. Schleissmann, K. Schleper, C. Schmitt, S. Schulze, A. Scott, E. Seabright, J. Shih, L. Sirot, S. Sita, M. Skuja, J. Stampfl, K. Stewart, W.C. Tucker, E. Urquhart, J. Vandermeer, K. van Atta, L. van Zuidam, J. Verge, G. Viallon, V. Vonau, R. Wakeford, A. Walker-Bolton, K. Watz, E. Wikberg, M. White, E. Williams, J. Williams, E. Wolf, D. Wood, D. Works, and M. Ziegler. Long-term site managers H. Gilkenson and W. Lammers made particularly large contributions.

### S1.2 Details about location data sampling

Fieldwork was conducted at the Lomas Barbudal Monkey Project, established by SEP in 1990, one of the longest-running longitudinal studies of white-faced capuchins and non-human primates more broadly. We used two types of location data: (1) sleep-site locations collected continuously from 1991–2023, and (2) handheld GPS data collected from 2009–2020, once the technology could reliably function under dense canopy. Full documentation on data processing, cleaning, and validation is available in Jacobson *et al.* [1]. Because only one or two groups could be monitored at a time, location sampling was irregular, with researchers following a given group for around 2-7 days on average to complete focal follows before switching groups. These sampling gaps were rigorously assessed and addressed in Jacobson *et al.* [2]. Sampling windows also varied by group, depending on when groups were first studied or formed via permanent fission from existing groups (see Figure S1).

### S1.3 Details about environmental data

#### S1.3.1 Quantifying habitat quality

We used surface reflectance data from Landsat 5, 7, and 8 to calculate NDVI (Normalized Difference Vegetation Index) from red (`RED`) and near-infrared (`NIR`) bands using the standard formula:

$$\mathrm{NDVI} = \frac{\mathrm{NIR} - \mathrm{RED}}{\mathrm{NIR} + \mathrm{RED}}$$

For each year and season, we generated pixel-wise maximum NDVI composites around the study site at a 30 m resolution. NDVI varied strongly across seasons and closely tracked the mean daily distance to rivers by all capuchin groups (Figure S2, Figure S3). This pattern reflects (a) the deciduous nature of the surrounding forest— where most trees lose their leaves during the dry season, especially away from permanent water sources— and (b) the monkeys' increased reliance on evergreen riparian zones during that time.

For a comprehensive synthesis of long-term environmental and climate variability in the Guanacaste region, we direct readers to Campos [4]. In that study, MODIS data were used to generate NDVI, resulting in substantially higher values than those observed in our Landsat-derived dataset. This discrepancy is likely due to MODIS's coarser spatial resolution (250 m), which can smooth over fine-scale heterogeneity in the landscape, such as canopy gaps, shadows, roads, pasturelands, and other anthropogenic or natural disturbances. To assess consistency, we retrieved MODIS NDVI data using the `MODISTools` `R` package [5] and directly compared it to Landsat NDVI for the 2015 wet and dry seasons. While absolute values differed— MODIS showed seasonal means of 0.45 (dry) and 0.85 (wet), compared to Landsat's 0.25 and 0.45— the magnitude of seasonal change was remarkably similar between sensors (Figure S4). We selected Landsat as our primary data source because its higher spatial resolution (30 m) allows for finer-scale estimation of habitat quality across capuchin monkey home ranges, and because it offers historical continuity back to the early 1970s (unlike MODIS, which begins in 2000), encompassing our long-term behavioral and demographic datasets. Campos [4] also compared NDVI with the Enhanced Vegetation Index (EVI), which is often considered more

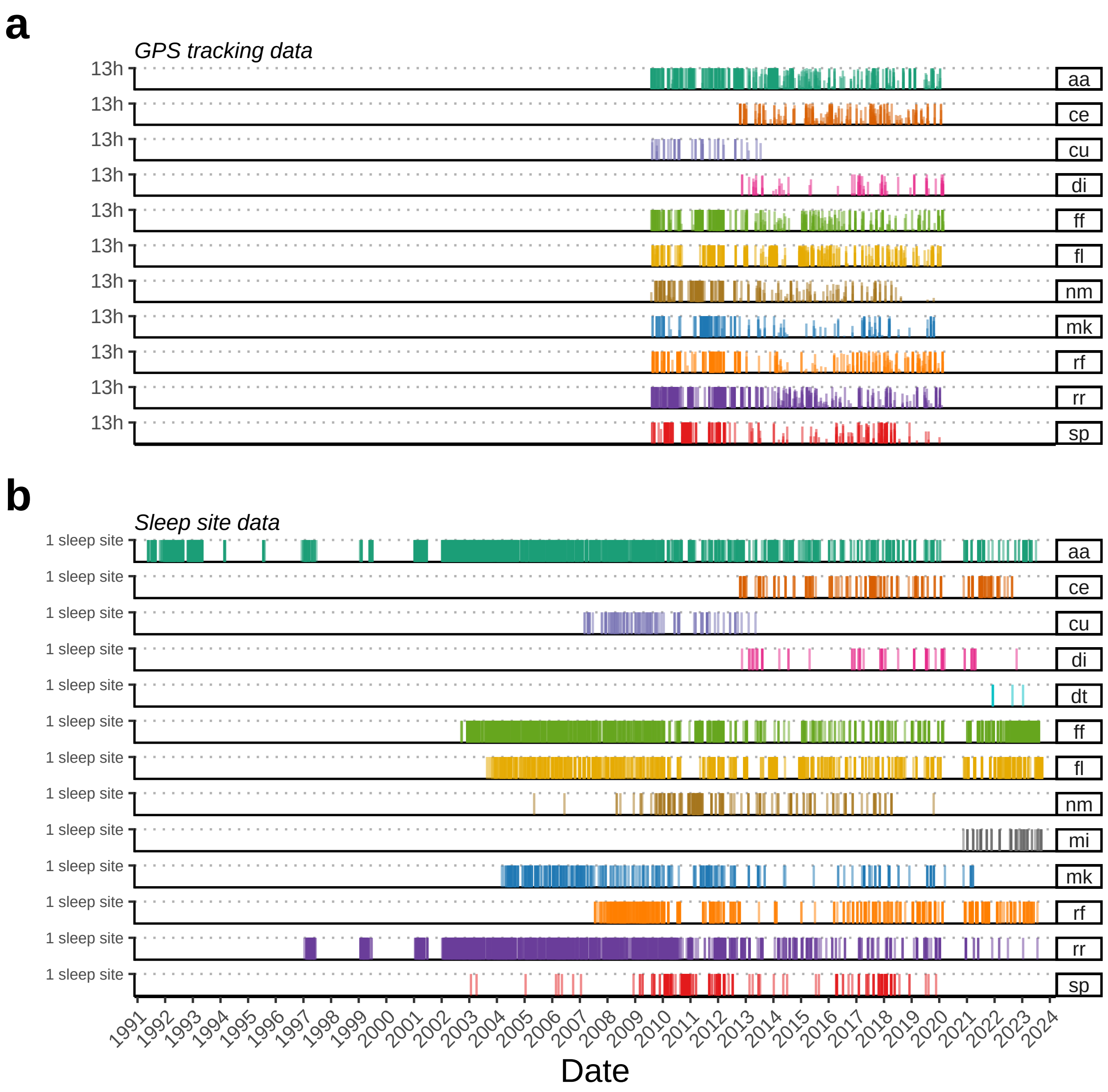


Figure S1: **Temporal distribution of location data collected from 13 groups of white-faced capuchins at the Lomas Barbudal Monkey Project, Costa Rica (1991–2023).** Panel (a) shows daily handheld GPS tracking effort, where each vertical line represents one day of observation (up to 13 tracking hours per day). Panel (b) shows sleep-site records, with each vertical line representing a single recorded sleep location. Lines are colored by group, with group names listed on the right. Figure generated using code adapted from Campos *et al.* [3].

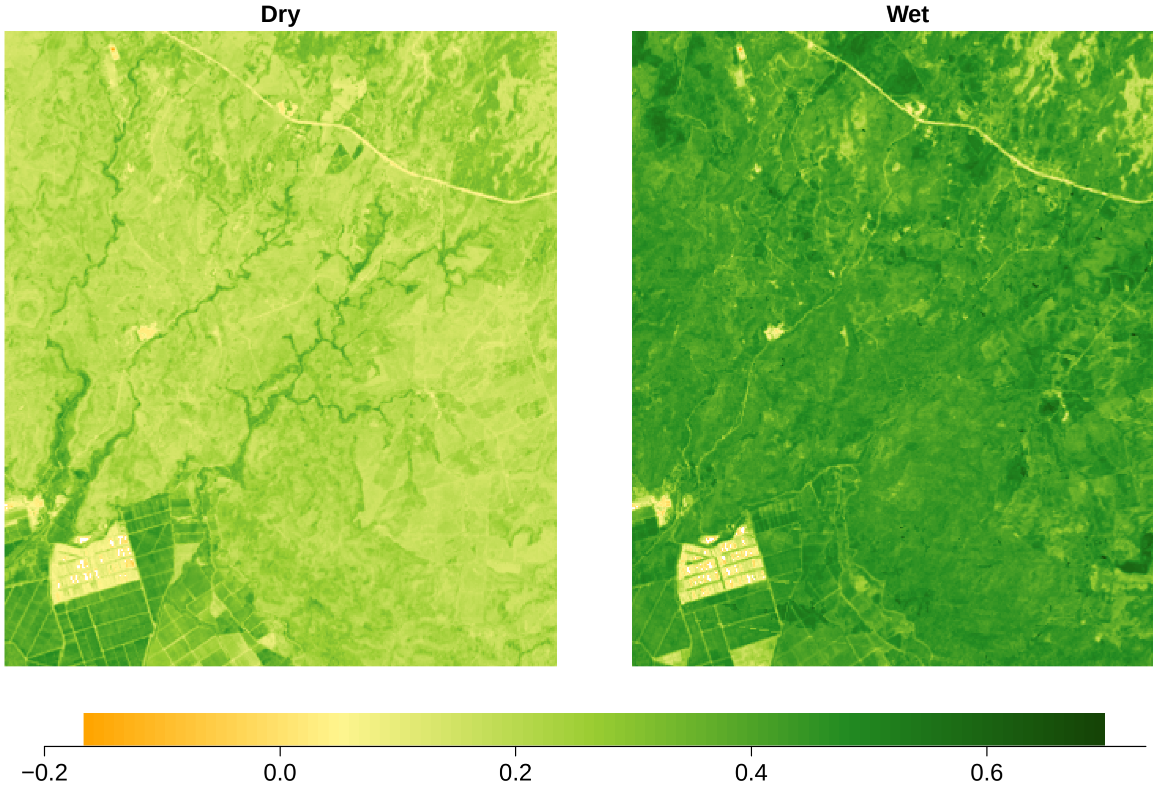


Figure S2: **Normalized Difference Vegetation Index (NDVI) seasonal composites for the Lomas Barbudal Monkey Project study area in 2015.** NDVI values were derived from Landsat 7, 8, and 9 surface reflectance imagery. Dry season composites include images from February to April; wet season composites include images from September to November. Pixel values represent the maximum NDVI observed across all images within each seasonal window, at 30 m resolution. Warmer colors (orange to yellow) indicate lower NDVI values, while darker greens represent higher NDVI values.

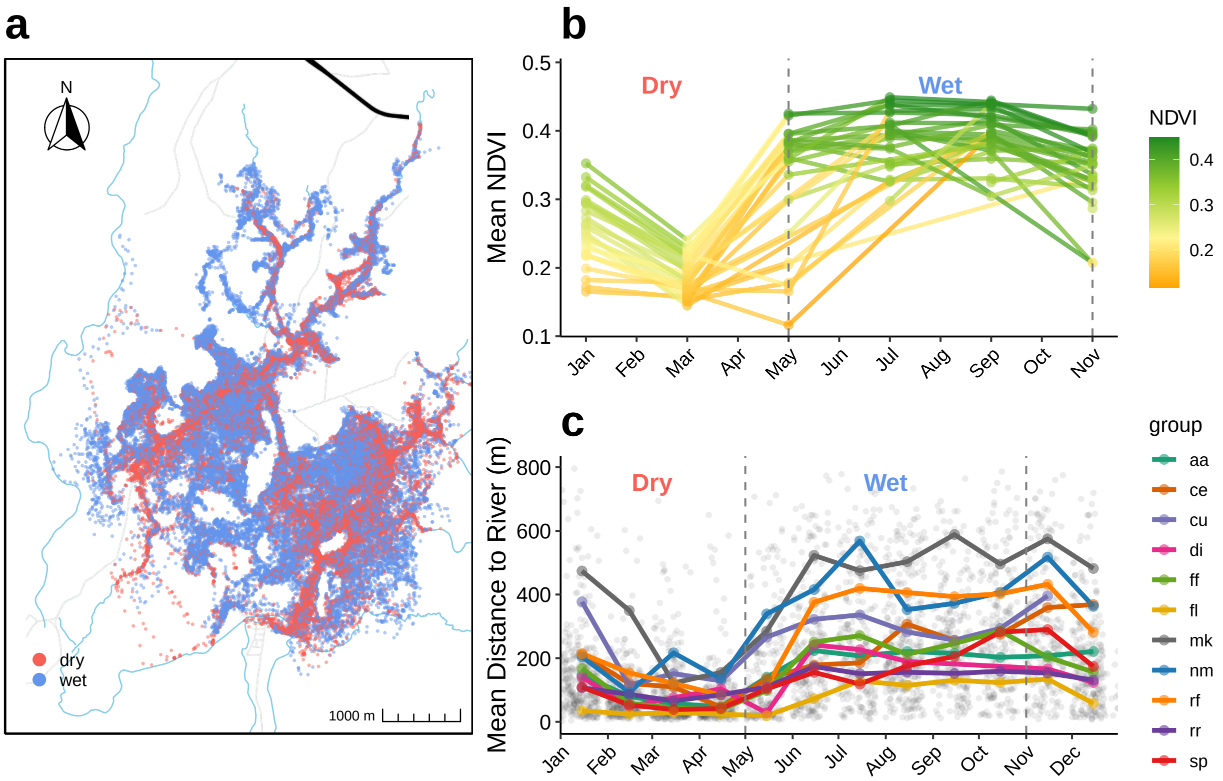


Figure S3: **Seasonal variation of Normalized Vegetation Index (NDVI) and capuchin locations relative to rivers at the Lomas Barbudal Monkeys Project.** (a) Handheld GPS-tracking data (30-minute fix rate) from all study groups, September 2009 - April 2020, colored by season (red = dry, blue = wet). (b) Bimonthly NDVI values from 1991 to 2023. Points represent landscape-scale means from bimonthly maximum-value composites; lines connect observations within the same year. Line colors correspond to the NDVI value at each observation (colors reflect the preceding point along each segment, not a continuous gradient along the line). Warmer colors (orange to yellow) indicate lower NDVI values, while darker greens represent higher values. Dashed vertical lines indicate typical transitions between dry and wet seasons. (c) Mean distance to rivers over time. Gray points show daily group-level means; colored points and lines represent monthly means per group.

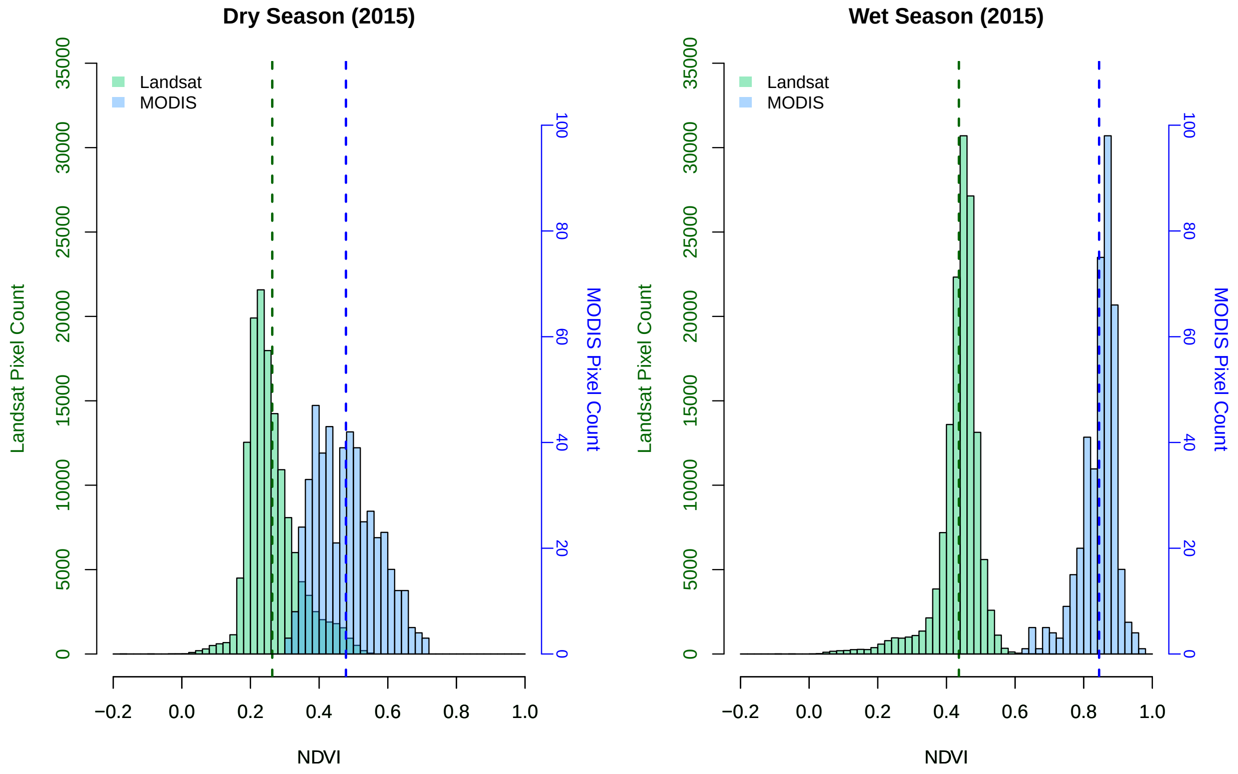


Figure S4: **Comparison of Landsat and MODIS NDVI distributions during the 2015 dry and wet seasons at the Lomas Barbudal Monkey Project.** Histograms show NDVI pixel values derived from Landsat (green, left y-axis) and MODIS (blue, right y-axis) for the dry (left panel) and wet (right panel) seasons. Dashed vertical lines indicate mean NDVI values for each sensor.

robust in tropical forests due to its reduced sensitivity to canopy saturation and atmospheric interference, and found the two indices to be highly correlated. We opted for NDVI due to its longer temporal availability, broader use in ecological research, and greater comparability across studies and time periods.

#### S1.3.2 Quantifying seasonal severity and its relationship with ENSO

Tropical dry forests, including Lomas Barbudal, are highly sensitive to global-scale climatic fluctuations driven by the El Niño–Southern Oscillation (ENSO) cycle [6]. ENSO alternates between abnormally warm, dry *El Niño* phases and cool, wet *La Niña* phases of varying intensity and duration [7, 8], both of which shift phenological cycles and thereby impact resource availability for animal populations [9]. El Niño events often bring droughts and wildfires that reduce canopy cover and water availability across the tropical Americas [10], increasing heat stress and altering animal physiology and behavior [11, 12], and reducing survival and reproductive success in some species [13]. In contrast, extreme La Niña periods can trigger floods and landslides and disrupt flowering and arthropod cycles, sometimes leading to famine and mass mortality

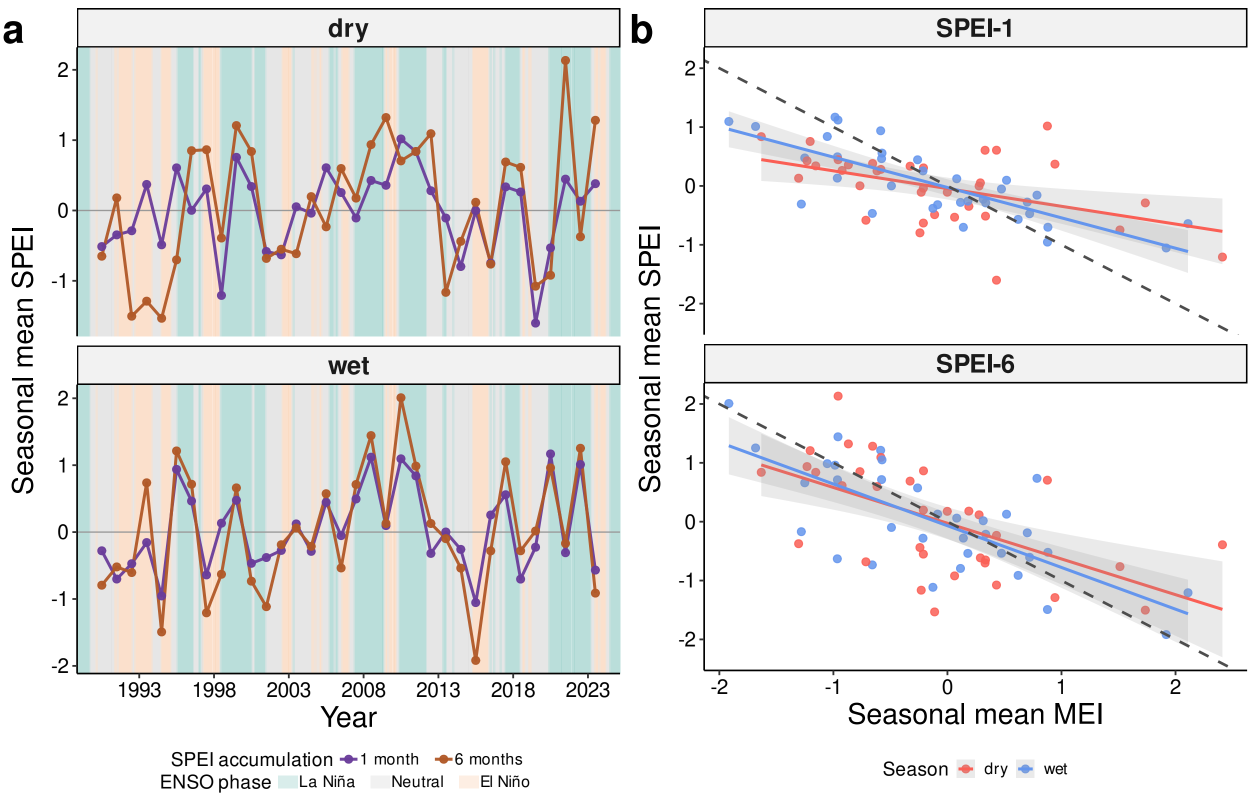


Figure S5: **Relationship between local hydroclimate (SPEI) and ENSO variability. (a)** Long-term trends in the Standardized Precipitation–Evapotranspiration Index (SPEI) from 1990–2023 around Lomas Barbudal Biological Reserve, calculated over 1-month (orange) and 6-month (purple) accumulation periods. Points show seasonal mean SPEI values per year, with lines connecting consecutive seasons. Negative SPEI values indicate abnormally dry periods and positive values indicate abnormally wet periods, relative to the typical range within each season (Jan–Apr = dry; May–Dec = wet). Shaded background bands denote phases of the El Niño Southern Oscillation (ENSO) cycle. **(b)** Relationship between seasonal mean SPEI and the Multivariate ENSO Index (MEI), a continuous measure of ENSO intensity (positive values = El Niño; negative = La Niña) obtained from the `rsoi` R package [18]. The dashed black line indicates a 1:–1 slope representing perfect negative correspondence between indices. Points represent seasonal means (red = dry, blue = wet); solid lines show fitted linear trends (±95% confidence intervals).

among mammals [14, 15]. Climate change is expected to intensify both phases of ENSO—making them more frequent, prolonged, and severe [16, 17]—which may exacerbate resource unpredictability and, in turn, reshape the balance of within- and between-group competition and the trade-offs of group-living.

We investigated how local hydroclimatic variation (i.e., seasonal severity) influences within- and between-group competition in the Lomas Barbudal capuchins, with the aim of generating predictions for how intensifying ENSO cycles may affect these dynamics under future climate change. Seasonal severity was quantified using the Standardized Precipitation–Evapotranspiration Index (SPEI), which measures anomalies in water balance calculated as precipitation minus potential evapotranspiration (PET)—the amount of water lost to evaporation and plant transpiration under existing temperature and radiation conditions. Monthly SPEI data (1990–2023) were obtained from the ERA5-Drought reanalysis dataset [19] for the Lomas Barbudal Biological Reserve and adjacent lands used by the study population. Values were standardized within seasons so that negative scores indicate abnormally dry and positive scores abnormally wet conditions relative to the typical range for that season (Jan–Apr = dry; May–Dec = wet). We compared SPEI calculated over 1-month and 6-month accumulation periods, representing short-term and long-term cumulative water balance, respectively. The 1-month SPEI reflects immediate within-season severity, whereas the 6-month SPEI (which partly integrates conditions from the preceding season) captures the broader climatic context and aligns more closely (though inversely) with global ENSO variability (Figure S5).

To evaluate how hydroclimatic anomalies modify competitive dynamics, we modeled the effects of group size across representative seasonal SPEI values (-1.5 = abnormally dry, 0 = average, 1.5 = abnormally wet). Mean home-range NDVI was used as a proxy for *between-group competitive ability*, as groups that more effectively defend or acquire high-quality areas are expected to occupy ranges with greater vegetation productivity. Per-capita fruit intake rate served as a proxy for *within-group competition*, as larger groups are predicted to experience reduced foraging efficiency due to scramble competition. Model details are provided in subsubsection 4.5.1.

The effects of group size on home-range NDVI were consistent between the 1-month and 6-month models: larger groups occupied higher-NDVI ranges during **dry seasons that were wetter than average** (SPEI $> 0$), but not during **abnormally dry dry seasons** (SPEI $< 0$) or the **wet season**, where relationships were negligible (though weakly positive when conditions were **drier than usual**). Thus, larger groups hold higher-quality ranges when **dry seasons are wetter than usual** (and possibly when **wet seasons are drier than usual**), regardless of timescale.

Differences between the two accumulation periods were more pronounced for fruit intake. Under short-term hydroclimate (SPEI-1), group size effects were weakly negative across all SPEI values in the **dry season**. Under longer-term hydroclimate (SPEI-6), this negative effect weakened (i.e., became more positive) from **exceptionally dry** to **exceptionally wet** conditions. In the **wet season**, group size had a more positive effect on fruit intake under **exceptionally dry conditions**, but this effect weakened or reversed as conditions **became wetter**—a pattern more pronounced in the short-term (SPEI-1) models. Thus, in the **dry season**, group-size effects are more responsive to longer-term moisture balance (SPEI-6), whereas in the **wet season**, they are more sensitive to short-term anomalies (SPEI-1).

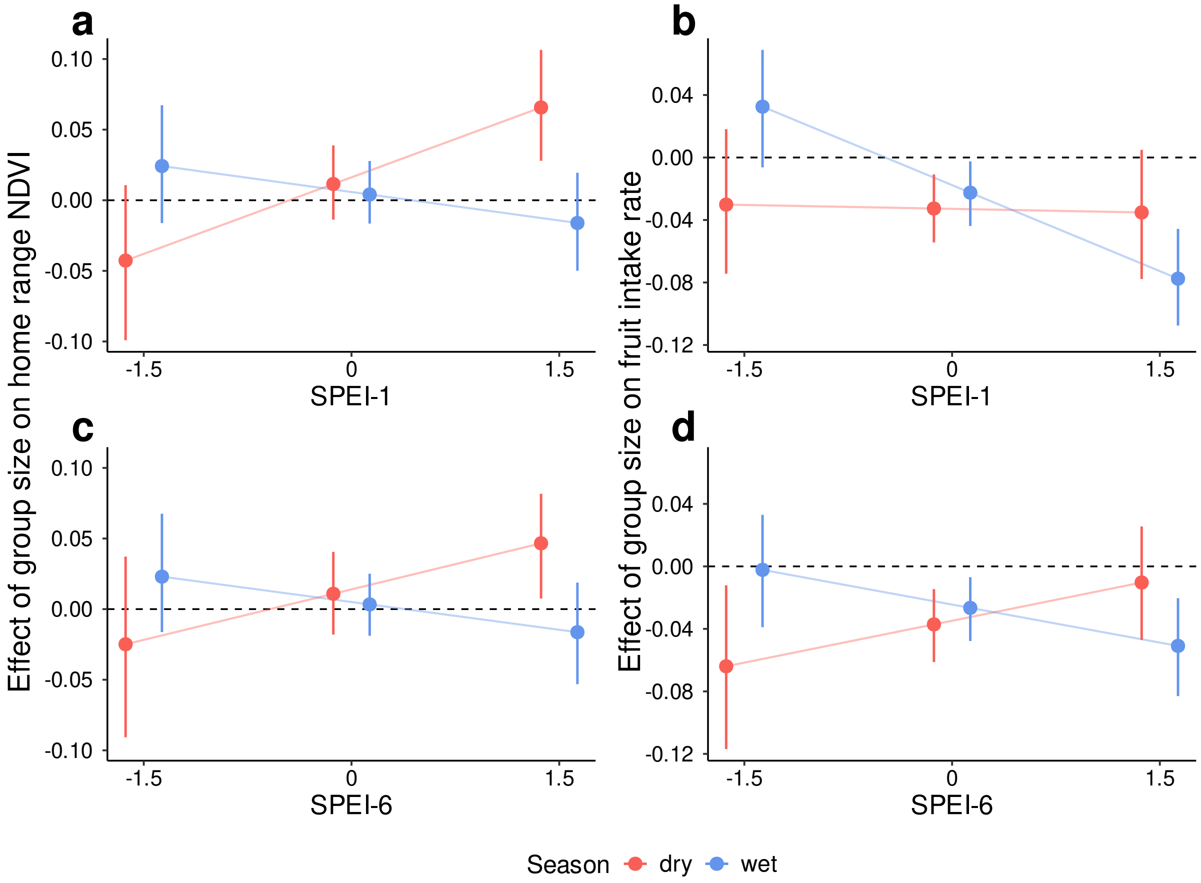


Figure S6: **Estimated effects of group size on within- and between-group competition across local hydroclimatic variation (SPEI).** Point intervals show posterior estimates of the effect of group size on mean home-range NDVI (panels **a**, **c**) and per-capita fruit intake rate (panels **b**, **d**) at representative SPEI values (–1.5, 0, 1.5), calculated using 1-month (**a**, **b**) and 6-month (**c**, **d**) accumulation periods. Points are posterior medians with 89% highest posterior density intervals (vertical bars); the dotted horizontal line indicates zero effect. Blue and red denote wet- and dry-season estimates, respectively. Light colored lines connect median effects across SPEI values to aid visual comparison. Negative SPEI values indicate abnormally dry periods and positive values indicate abnormally wet periods, relative to the typical range within each season (Jan–Apr = dry; May–Dec = wet). Models used $n = 224$ seasonal home ranges across 12 groups.

### S1.4 Details about causal assumptions

To determine appropriate covariate structure for estimating our response variables and permit unbiased estimation of causal effects, we used the `dagitty` package to construct directed acyclical graphs or "DAGs" [20]. DAGs provide a visual framework for representing hypothesized causal relationships among variables in a given model or research question. Based on our a priori assumptions about the biological system, we identified adjustment sets using the `adjustmentSets` function, which returns covariate combinations that— if our causal assumptions are correct— permit unbiased estimation by statistically conditioning for confounding variables (i.e., back-door paths).

#### S1.4.1 Assumptions for estimating daily path length, revisitation rate, home range area and quality

Below are our assumptions regarding the causal pathways for the GLMMs estimating (a) per capita fruit intake rate, (b) daily path length, (c) revisitation rate, (d) home range area, and (e) home range quality as depicted in the DAGs shown in Figure S7, Figure S8, Figure S9, and Figure S10. Habitat quality is measured via the mean Normalized Vegetation Index (NDVI) within a group's annual or seasonal home range boundary. We define revisitation as both the frequency with which groups return to areas across their range and the extent of their exploratory behavior. We consider these to be two sides of the same coin, and therefore use revisitation rate and exploration interchangeably when discussing the causal assumptions below.

1. Season → Habitat Quality: During the dry season, deciduous trees, which comprise 70-80% of the forest at Lomas Barbudal, shed their leaves, reducing canopy cover and concentrating fruiting trees and other key resources near evergreen riparian areas [21].

2. Season → Daily Path Length and Home Range Area: Seasonal climate variation affects both variables directly— via heat stress and physiological limits— and indirectly— by altering habitat quality and resource distribution.

3. Habitat Quality → Fitness: Vegetation quality, distribution, and abundance affect access to key resources that are critical for individual survival and reproduction.

4. Fitness → Group Size: The number of individuals in a group is determined by individual survival and reproduction.

5. Group Size → Fruit Intake: Due to within-group competition, larger groups have decreased per capita fruit foraging efficiency.

6. Group Size → Competitive Power: Larger groups can more likely win intergroup contests and displace smaller groups.

7. Fruit Intake → Daily Path Length and Revisitation: Per capita fruit ingestion rate influences patch depletion rate, which in turn affects how far groups travel to find new resources and how often they revisit specific patches.

8. Fruit Intake → Fitness: Per capita fruit ingestion rate influences energy available for survival and reproduction.

9. Competitive Power → Revisitation: More dominant groups will explore further out beyond their core area resulting in fewer revisits to specific patches.

10. Revisitation → Home Range Area: Exploratory behavior and recursive movement patterns influence the extent to which groups move, thereby shaping their home range boundaries.

11. Age and Sex → Fruit Intake: Capuchins of different ages and sexes have varying metabolic requirements, which influence their fruit intake rates.

In this framework, we explicitly represent group size as a proxy for both between-group competition (via competitive power) and within-group competition (via per capita fruit intake rate). This approach underscores the inherent difficulty in isolating the causal effects of these different forms of competition on our outcomes of interest. Assuming our DAG is correct, exploratory movement and home range size are shaped by both forms of competition: larger groups may range more widely either due to increased resource needs (within-group) or greater ability to displace neighbors (between-group). Similarly, while within-group competition directly affects fruit intake rate, between-group competition may also influence habitat quality, which in turn impacts fruit intake— further linking the two processes through shared pathways. These overlapping mechanisms, combined with mismatched measurement scales— e.g., individual vs. group level, daily vs. seasonal— complicate proper adjustment for confounders. Accordingly, the models in this section should be interpreted as predictive rather than causal. This motivates our use of the Social Relations Model to estimate proportional overlap between neighbors $\mathrm{PO}_{fn}$ (see section 4.4) as a way to isolate the influence of between-group competition.

#### S1.4.2 Assumptions for estimating proportional overlap, $\mathrm{PO}_{fn}$

Below are our assumptions regarding the causal pathways for the Social Relations Model estimating proportional overlap $\mathrm{PO}_{fn}$, as depicted in the DAG shown in Figure S11. We do not repeat assumptions that are

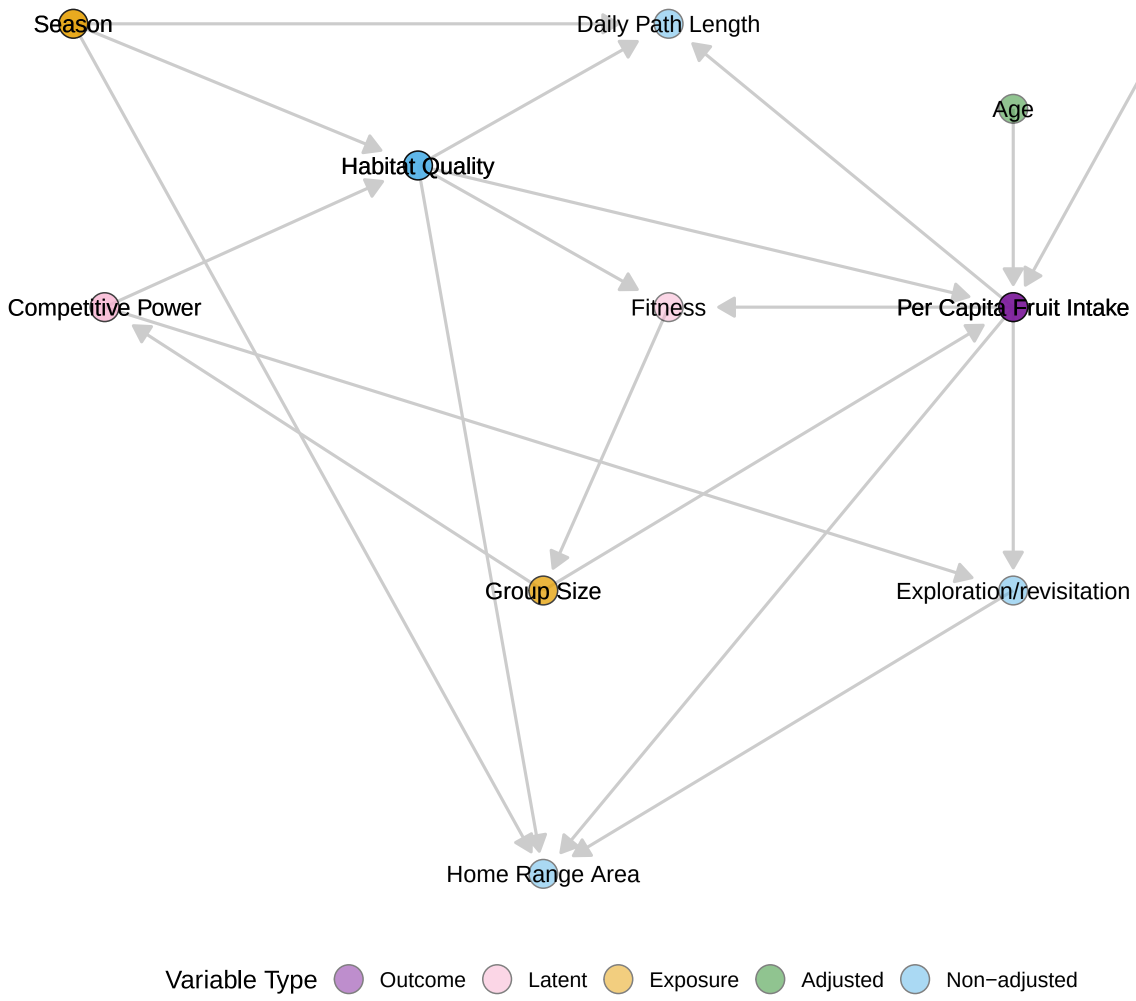


Figure S7: **Directed Acyclic Graph (DAG) illustrating proposed causal pathways affecting the outcome of interest: per capita fruit intake rate.** The outcome variables are shown in purple, which were estimated in separate models but with the same covariate structure. Orange nodes are predictors of interest (i.e., exposure variables). Pink nodes represent latent (unobserved) variables, green nodes are covariates adjusted for to block back-door paths, and blue nodes are variables not adjusted for because adjustment was unnecessary. Arrows indicate proposed causal directions.

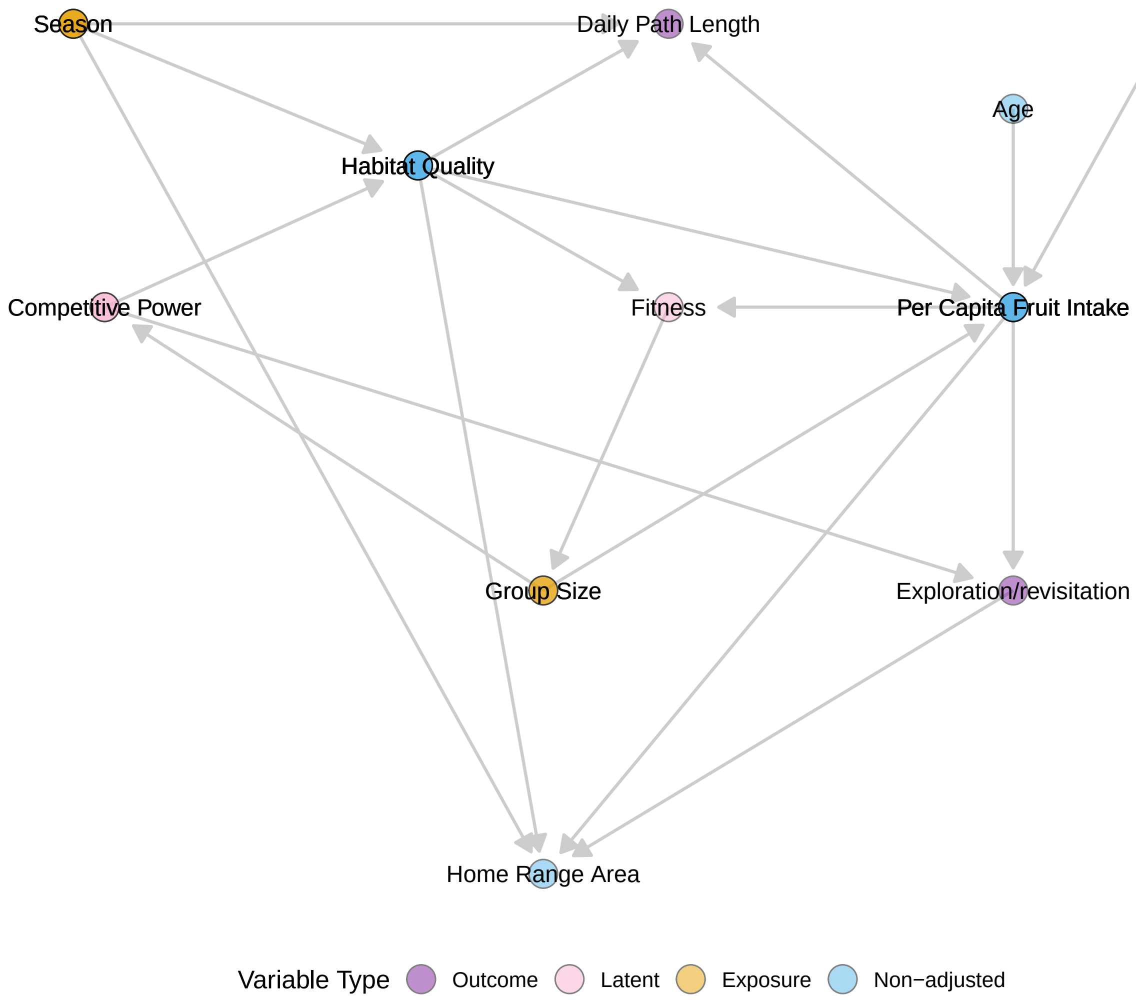


Figure S8: **Directed Acyclic Graph (DAG) illustrating proposed causal pathways affecting the outcomes of interest: daily path length and revisitation rate.** The outcome variables are shown in purple, which were estimated in separate models but with the same covariate structure. Orange nodes are predictors of interest (i.e., exposure variables). Pink nodes represent latent (unobserved) variables, green nodes are covariates adjusted for to block back-door paths, and blue nodes are variables not adjusted for because adjustment was unnecessary. Arrows indicate proposed causal directions.

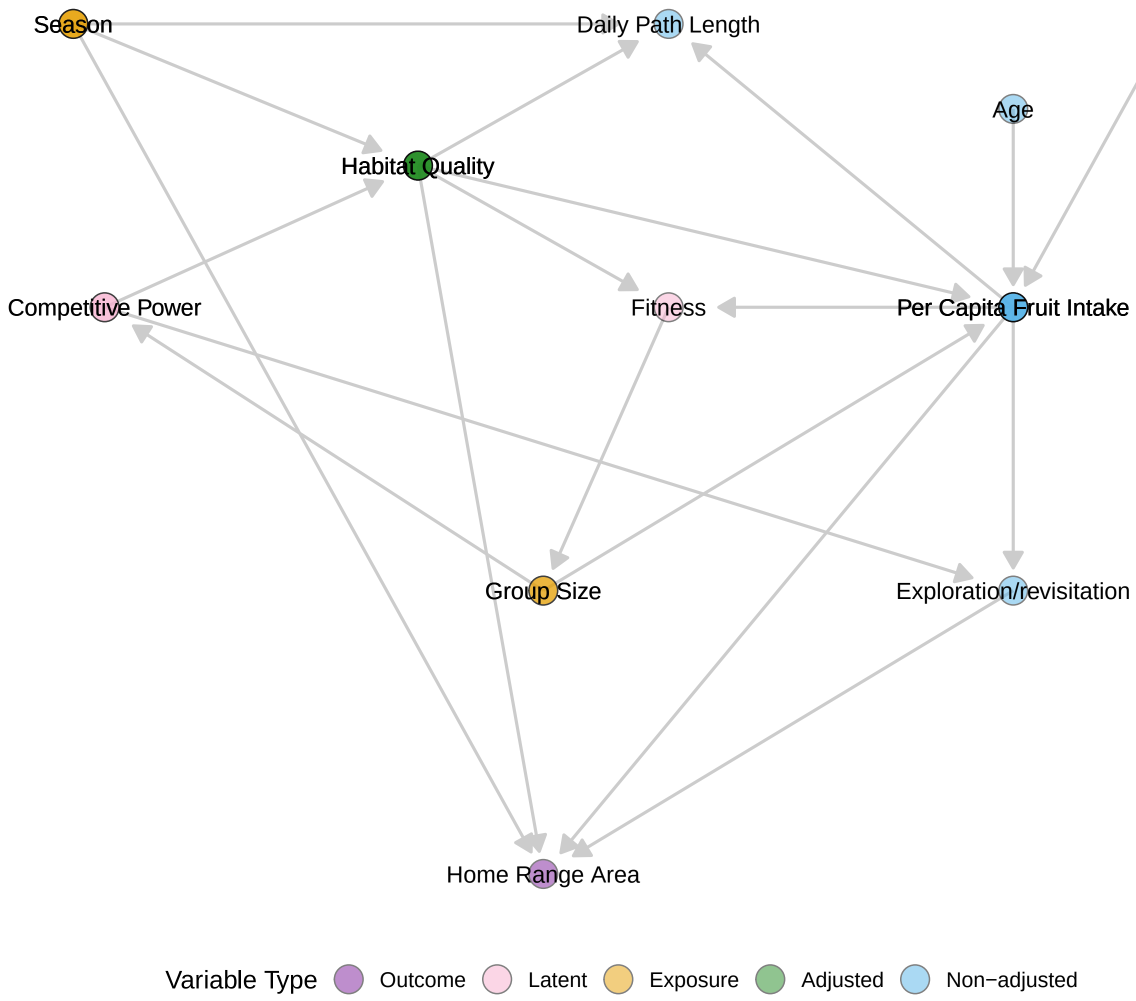


Figure S9: **Directed Acyclic Graph (DAG) illustrating proposed causal pathways affecting the outcome of interest: home range area.** The outcome variable is shown in purple. Orange nodes are predictors of interest (i.e., exposure variables). Pink nodes represent latent (unobserved) variables, green nodes are covariates adjusted for to block back-door paths, and blue nodes are variables not adjusted for because adjustment was unnecessary. Arrows indicate proposed causal directions.

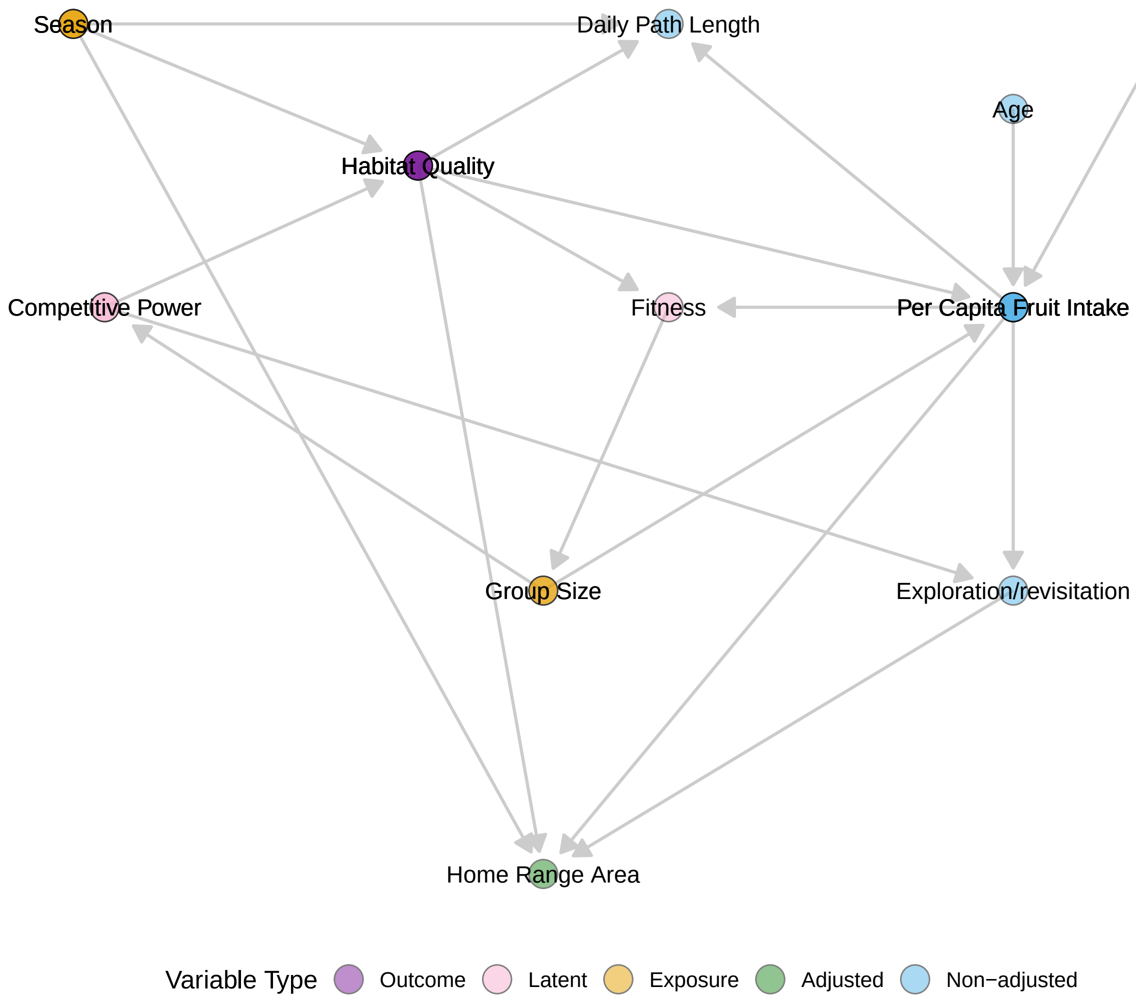


Figure S10: **Directed Acyclic Graph (DAG) illustrating proposed causal pathways affecting the outcome of interest: home range quality.** The outcome variable is shown in purple. Orange nodes are predictors of interest (i.e., exposure variables). Pink nodes represent latent (unobserved) variables, green nodes are covariates adjusted for to block back-door paths, and blue nodes are variables not adjusted for because adjustment was unnecessary. Arrows indicate proposed causal directions.

identical to those listed in the previous section.

1. Focal Group Size → Proportion Overlap: Larger focal groups, due to their numerical advantage and stronger competitive ability, can encroach further into the ranges of smaller neighboring groups.

2. Neighbor Group Size → Proportion Overlap: Larger neighboring groups, due to their numerical advantage and stronger competitive ability, can encroach further into the ranges of smaller focal groups.

3. Focal Home Range Area → Proportion Overlap: Given a constant intersection area, a larger focal home range will have less proportion overlap. This does not apply for neighbor home range area because proportion overlap is measured from the focal group's perspective.

4. Neighbor? (Y/N) → Proportion Overlap: Whether or not the two groups are neighbors determines whether the proportion overlap is zero or non-zero.

Note that we assumed a causal relationship between focal home range area (HRA) and $\text{PO}_{fn}$, but not neighbor HRA and $\text{PO}_{fn}$. This is because focal HRA appears directly in the denominator when calculating $\text{PO}_{fn}$ (see section 4.4). Given a constant intersection area, a larger focal home range will naturally result in a lower proportional overlap. Since our goal is to estimate the degree of encroachment by the neighbor group within each dyad, it is essential to adjust for focal HRA to control for this structural dependency and avoid confounding bias. In contrast, the same reasoning does not apply to neighbor HRA. Adjusting for it would, in fact, obscure the effect of neighbor group size on $\text{PO}_{fn}$, as it could absorb part of the variation we aim to attribute to neighboring group behavior. For this reason, we do not treat neighbor HRA as having a direct causal effect on $\text{PO}_{fn}$, unlike focal HRA.

The following adjustment sets were considered valid for estimating the direct effects of focal group size, neighbor group size, and season on $\text{PO}_{fn}$:

1. $\{\text{Focal HRA}, \text{Neighbor? (Y/N)}\}$

2. $\{\text{Focal HRA}, \text{Focal Habitat Quality}, \text{Neighbor Habitat Quality}\}$

We selected the first adjustment set because satellite imagery required to calculate NDVI-based habitat quality was limited before 2000. Using the second set would have forced us to trim our spatial and demographic datasets to match the availability of those images.

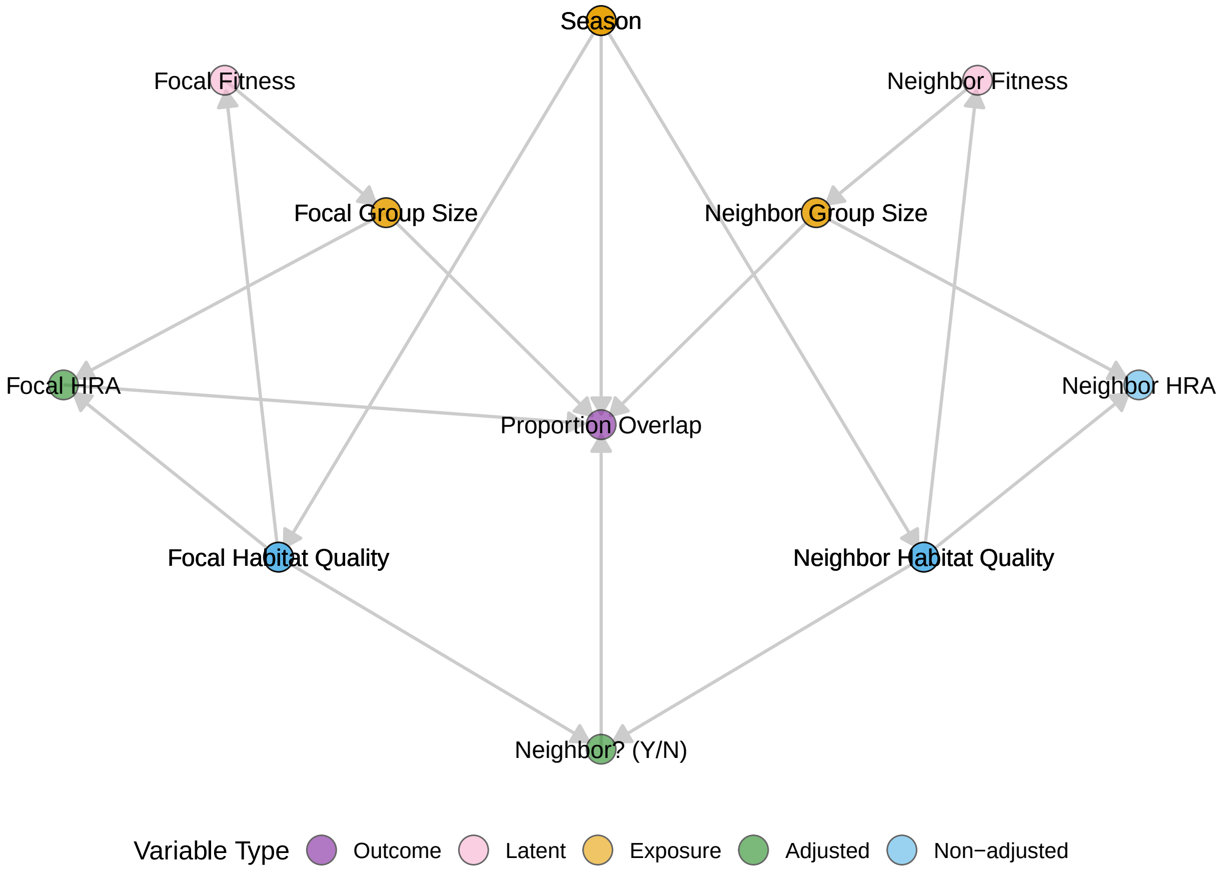


Figure S11: **Directed Acyclic Graph (DAG) illustrating proposed causal pathways affecting the outcome of interest: the proportion of the focal group's home range that overlaps with a particular neighbor, $\mathbf{PO}_{fn}$.** The outcome variable is shown in purple. Orange nodes are predictors of interest (i.e., exposure variables). Pink nodes represent latent (unobserved) variables, green nodes are covariates adjusted for to block back-door paths, and blue nodes are variables not adjusted for because adjustment was unnecessary. Arrows indicate proposed causal directions.

#### S1.4.3 Assumptions for estimating encounter rates

Below are the assumed causal pathways informing our model of intergroup encounter rates, as depicted in the DAG in Figure S12. We do not repeat assumptions already stated in previous sections.

1. Focal and Neighbor Group Size → Group Size Difference: Defined as the absolute difference between focal and neighbor group sizes, representing asymmetry in competitive ability.

2. Group Size Difference → Encounter Rate: Greater asymmetry in group size may influence encounter frequency due to differences in competitive ability.

3. Neighbor (Y/N)? → Encounter Rate: Only neighboring groups share spatial proximity, making encounters possible.

4. Season → Encounter Rate: Seasonal variation affects encounter rates both indirectly—by concentrating resources (e.g., in riparian zones during the dry season)—and directly, via altered movement patterns due to temperature and water availability.

5. Overlap Area → Encounter Rate: Increased spatial overlap between home ranges raises the likelihood of intergroup encounters.

6. Overlap Habitat Quality → Encounter Rate: Higher-quality habitat in overlap zones may increase usage, thereby increasing encounter probability.

Based on the DAG structure, the only valid adjustment set for estimating the direct effects of group size difference, overlap habitat quality, and season on encounter rate was: $\{\text{Overlap Area}, \text{Neighbor (Y/N)}\}$. We adjusted for these covariates accordingly.

### S1.5 Details about Social Relations Model

To investigate the effect of relative group size on proportional overlap $PO_{fn}$, we used an extension of the multilevel Social Relations Model (SRM) [22]. Specifically, we implemented a zero-augmented (hurdle) SRM that distinguishes between (1) the probability of any overlap occurring (modeled with a Bernoulli distribution) and (2) the extent of overlap when present (modeled with a Beta distribution). Relative group size was modeled using focal group size, neighbor group size, and their interaction as fixed effects. Seasonality was incorporated as a three-way interaction between season, focal group size, and neighbor group size. To account for non-independence within dyads, we included varying effects for focal group ID, neighbor group

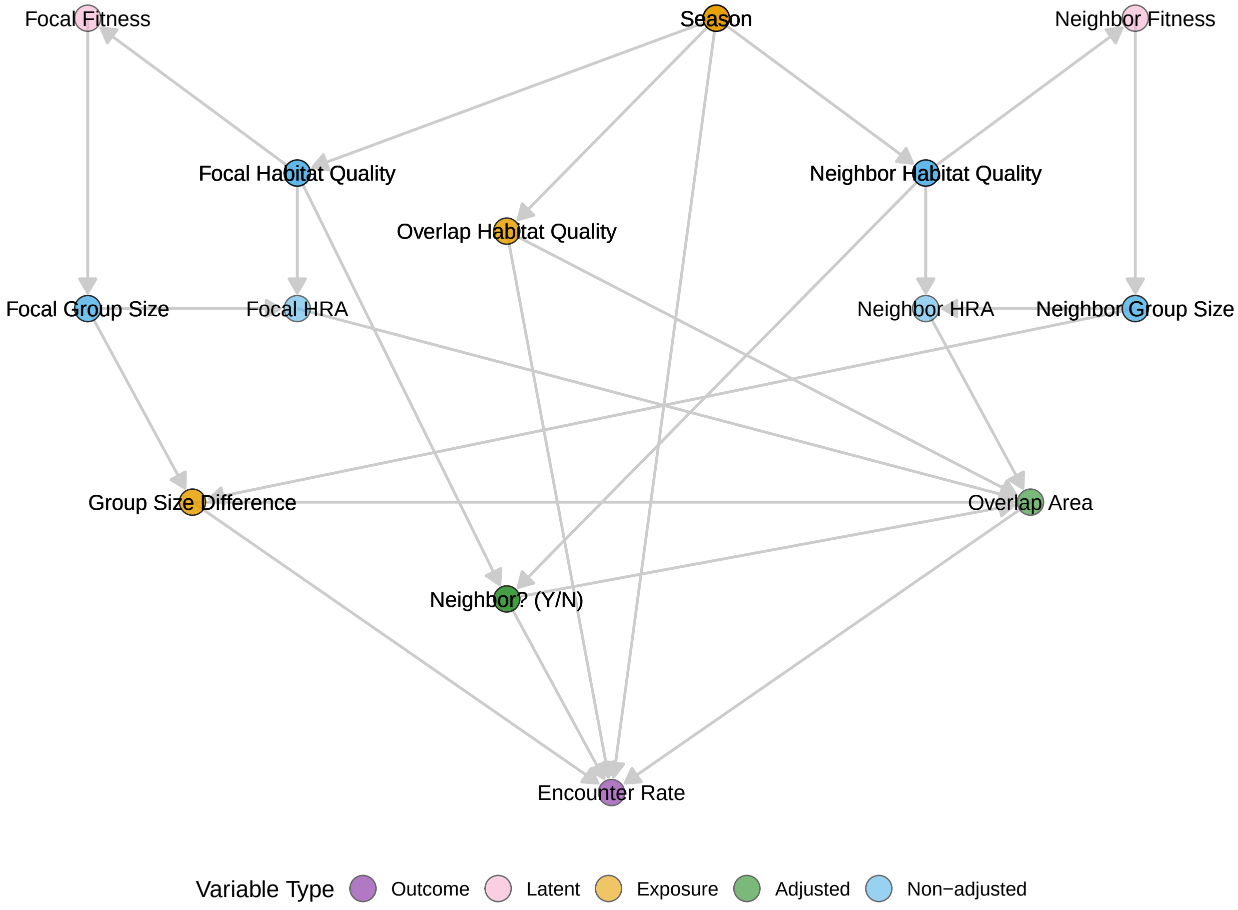


Figure S12: **Directed Acyclic Graph (DAG) illustrating proposed causal pathways affecting the outcome of interest: intergroup encounter rate.** The outcome variable is shown in purple. Orange nodes are predictors of interest (i.e., exposure variables). Pink nodes represent latent (unobserved) variables, green nodes are covariates adjusted for to block back-door paths, and blue nodes are variables not adjusted for because adjustment was unnecessary. Arrows indicate proposed causal directions.

ID, and dyad ID, modeled jointly via a multivariate distribution with a shared correlation structure. The full statistical specification of the model is provided below:

The probability of non-zero overlap between focal group $i$ and neighbor group $j$ at time $t$ is specified as:

$$Z_{|ij|,t} \sim \text{Bernoulli}(p_{|ij|,t})$$

$$\text{logit}(p_{|ij|,t}) = \bar{\alpha_0} + f_{0,i} + n_{0,j} + (\beta\bar{D}_0 + \beta D_{0,|ij|}) \cdot distance_t$$

Here, $\bar{\alpha_0}$ is the mean intercept across all groups and dyads, $f_i$ is a group-level varying intercept for focal group $i$, $n_j$ is a group-level varying intercept for neighbor group $j$, $\beta\bar{D}_0$ is the overall fixed effect quantifying the distance between home range centroids across all dyads, and $\beta_0 D_{|ij|,t}$ is the dyad-level varying fixed effect quantifying the distance between home range centroids of groups $i$ and $j$.

The proportion of overlap with neighbor $j$ within the home range of focal group $i$ at time $t$ is specified as:

$$P_{\vec{ij},t} \sim \text{Beta}(o_{\vec{ij},t}, \theta)$$

$$\text{logit}(o_{\vec{ij},t}) = \alpha_1 + \beta G_f \cdot \text{gs}_{f,t} + \beta G_n \cdot \text{gs}_{n,t} + \beta S \cdot \text{season}_t + \beta GS_{fns} \cdot \text{gs}_{n,t} \cdot \text{gs}_{f,t} \cdot \text{season}_t + \beta H_f \cdot \text{hr}_{f,t}$$

$$\alpha_1 = \bar{\alpha_1} + f_{1,i} + n_{1,j} + d_{1,|ij|}$$

$$\beta G_f = \beta\bar{G}_f + \beta G_{f,i}$$

$$\beta G_n = \beta\bar{G}_n + \beta G_{n,j}$$

$$\beta GS_{fns} = \beta G\bar{S}_{fns} + \beta GS_{fns,i} + \beta GS_{fns,j} + \beta GS_{fns,|ij|}$$

$$\beta H_f = \beta\bar{H}_f + \beta H_{f,i}$$

where $\alpha_1$ is the intercept, $\beta G_f$ is the fixed effect for focal group size, $\beta G_n$ is the fixed effect for neighbor group size, $\beta S$ is the fixed effect for season, $\beta GS_{fns}$ is the fixed effect for the interaction between focal group size, neighbor group size, and season, and $\beta H_f$ is the fixed effect for the home range area of the focal group. These parameters are further specified using submodels, where $\bar{\alpha_1}$ is the overall intercept, $f_{1,i}$ is a group-level varying intercept for focal group $i$, $n_{1,j}$ is a group-level varying intercept for neighbor group $j$, $d_{1,|ij|}$ is a dyad-level varying intercept, $\beta\bar{G}_f$ is the overall fixed effect for focal group size, $\beta G_{f,i}$ is the group-level varying slope for the group size of focal group $i$, $\beta\bar{G}_n$ is the overall fixed effect for neighbor group size, $\beta G_{n,j}$ is the group-level varying slope for the group size of neighbor group $j$, $\beta G\bar{S}_{fns}$ is the overall fixed effect for the interaction term, $\beta GS_{fns,i}$ is the group-level varying slope for focal group $i$ on the interaction, $\beta GS_{fns,j}$ is the group-level varying slope for neighbor group $j$ on the interaction, $\beta GS_{fns,|ij|}$ is the dyad-level varying slope on the interaction, $\beta\bar{H}_f$ is the overall fixed effect for focal home range area, and $\beta H_{f,i}$ is the group-level varying slope for the home range area of focal group $i$.

The final joint likelihood of the model is the product of the likelihoods from the Bernoulli and Beta components:

$$PO_{\vec{fn}} = \prod_{i,j,t} \left[\Pr(Z_{|ij|,t} = 1) \cdot f(P_{\vec{ij},t} \mid Z_{|ij|,t} = 1)\right]^{Z_{|ij|,t}} \cdot \left[\Pr(Z_{|ij|,t} = 0)\right]^{1-Z_{|ij|,t}} \quad (4)$$

where:

- $\Pr(Z_{|ij|,t} = 0)$ is the probability of observing zero overlap,
- $\Pr(Z_{|ij|,t} = 1)$ is the probability of observing non-zero overlap,
- $f(P_{\vec{ij},t} \mid Z_{|ij|,t} = 1)$ is the likelihood of the observed proportion of overlap, conditional on the overlap being non-zero.

### S1.6 Group-level varying effects

In most of our models, we included varying slopes and intercepts by group, allowing for group-specific predictions. These predictions enable us to examine how the relationship between group size and behavioral variables may differ across groups in subtle but important ways. Some groups inhabit especially fragmented landscapes due to roads and pasturelands, which affects their ranging patterns and habitat use (e.g., home range crossing time; [2]). These ecological differences can result in distinctive patterns for specific groups—most notably group FL.

FL's home range is highly fragmented, consisting of narrow riparian corridors winding through pasturelands, creating a patchy, forked canopy structure and generally lower habitat quality than other groups (Table S1). Unlike most other study groups, FL overlaps minimally with neighbors, including non-study groups, and occupies a largely exclusive range (Figure S13; MI, which split from FL in 2021 and occupies similar habitat, also has little contact with other groups, but appears to have moderate total overlap because much of its range is shared with FL; Figure 1). These ecological constraints can manifest in several ways. For example, while group size showed little to no effect on daily path length (DPL) at the population level and in most groups, it had a notably negative effect for group FL (Figure S14). This may reflect the increased risk or energetic costs of traversing a fragmented landscape— particularly one that requires frequent road or

Table S1: **Posterior predicted differences in home range quality (mean NDVI) between `fl` and other groups.** The final column shows the posterior probability that `fl` has lower NDVI.

| Group comparison | Mean diff [89% HPDI] | P(fl < other) |
|---|---|---|
| fl_vs_aa | -0.008 [-0.015, 0.000] | 0.960 |
| fl_vs_ce | -0.006 [-0.014, 0.002] | 0.890 |
| fl_vs_cu | -0.007 [-0.015, 0.002] | 0.860 |
| fl_vs_di | -0.005 [-0.014, 0.003] | 0.850 |
| fl_vs_ff | -0.005 [-0.014, 0.001] | 0.840 |
| fl_vs_mi | -0.001 [-0.011, 0.005] | 0.600 |
| fl_vs_mk | -0.006 [-0.013, 0.002] | 0.850 |
| fl_vs_nm | -0.004 [-0.011, 0.003] | 0.800 |
| fl_vs_rf | -0.003 [-0.009, 0.004] | 0.750 |
| fl_vs_rr | -0.007 [-0.016, 0.002] | 0.880 |
| fl_vs_sp | -0.001 [-0.008, 0.008] | 0.560 |

pasture crossings [23, 24]— or the added challenge of maintaining coordination in a large group navigating such an interrupted, non-contiguous environment [25]. An alternative possibility is historical: FL's negative group size–DPL relationship may partly reflect abnormally long path lengths during the period immediately after they split from AA group, when they were still a relatively small group attempting to establish a home range.

While the relationship between group size and DPL varied subtly across groups, the association between group size and home range area was consistently positive (Figure S15). To assess how ecological constraints impact space use, most prior studies have relied on between-group or cross-species comparisons [3, 26, 27], which are often confounded by geographic variation in habitat quality [28]. In contrast, our findings offer rare within-group evidence that home range area expands with group size over time— helping to disentangle demographic effects from environmental ones. As noted in Jacobson *et al.* [1], this kind of long-term, within-group analysis is essential for testing key predictions of the Ecological Constraints Model.

With respect to our findings on dyadic proportional overlap ($PO_{fn}$), we observed notable group-level variation. Specifically, by comparing each group's general tendency to give overlap to neighbors (neighbor group-level varying intercept) with their tendency to receive overlap from neighbors (focal group-level varying intercept), we find that groups that remained relatively small throughout the study tend to receive markedly more overlap than they give (Figure S16). In contrast, groups with larger average sizes and/or greater variability in group size over time exhibit stronger positive correlations between generalized giving and receiving of overlap. These findings support the idea that larger groups leverage their numerical advantage to expand into the ranges of their smaller neighbors.

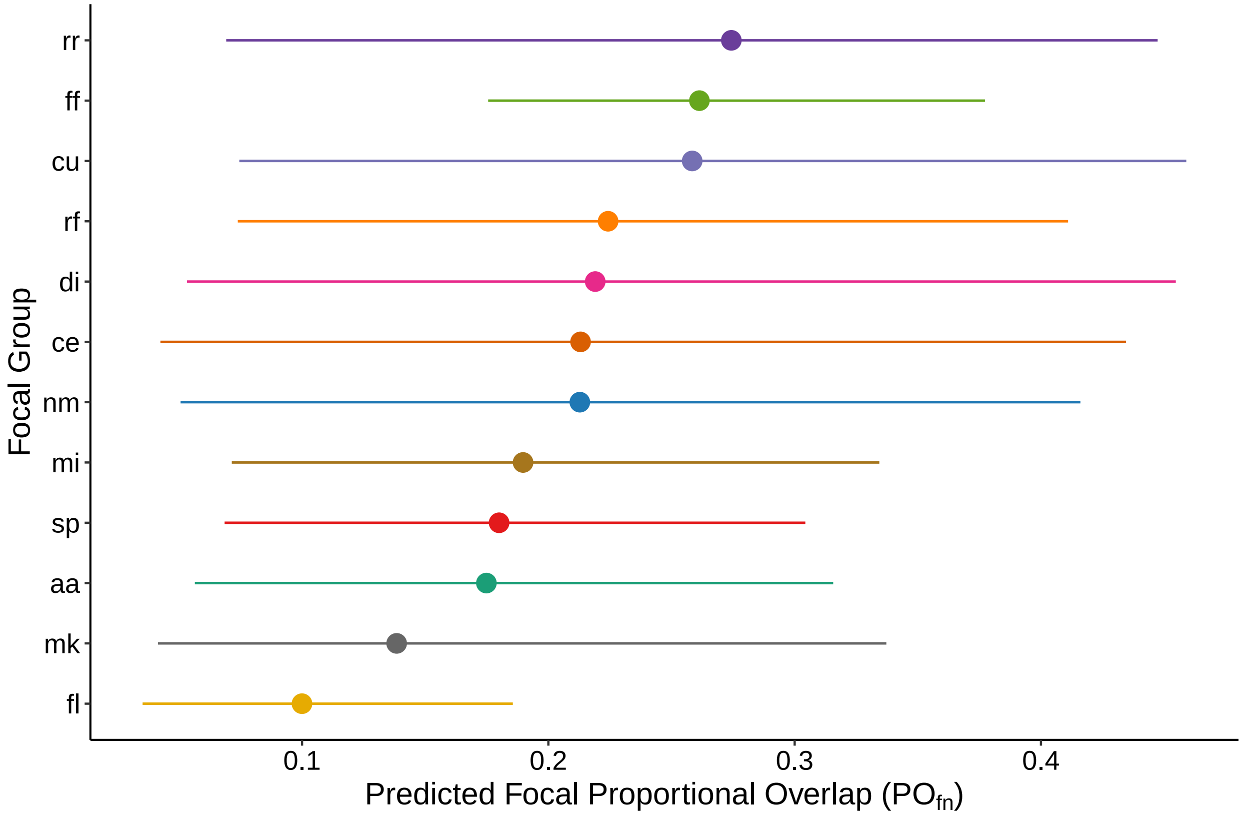


Figure S13: **Group-level model predictions for baseline proportional overlap ($PO_{fn}$) across groups.** Colors correspond to group identity. Points represent posterior medians and lines show 89% Highest Posterior Density Intervals. Model was fit to $n = 930$ dyad-years (59 dyads, 12 groups).

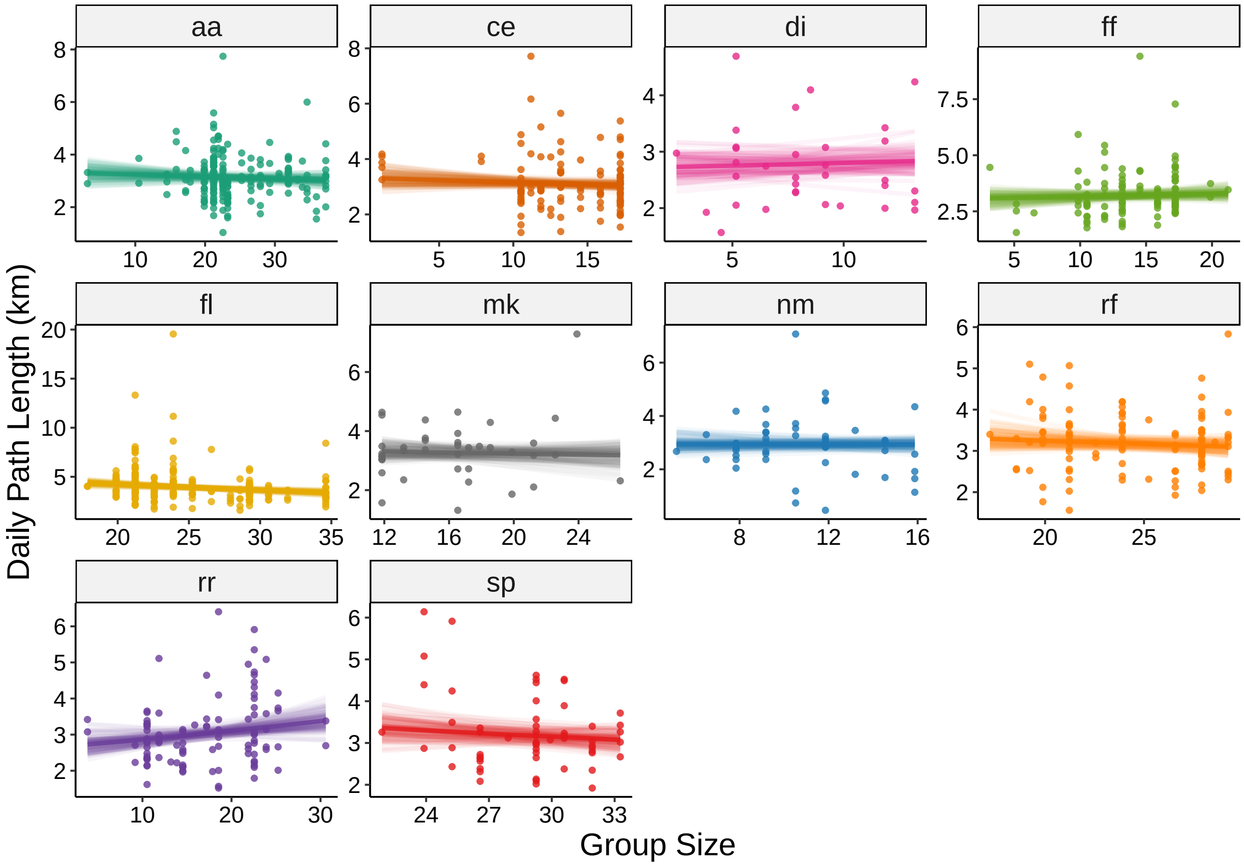


Figure S14: **Group-level model predictions for changes in daily path length (DPL) as a function of group size.** Colors correspond to group identity. Lighter lines are 100 randomly sampled posterior predictions, with the posterior median as a dark line. Model was fit to $n = 996$ group-days across 11 groups.

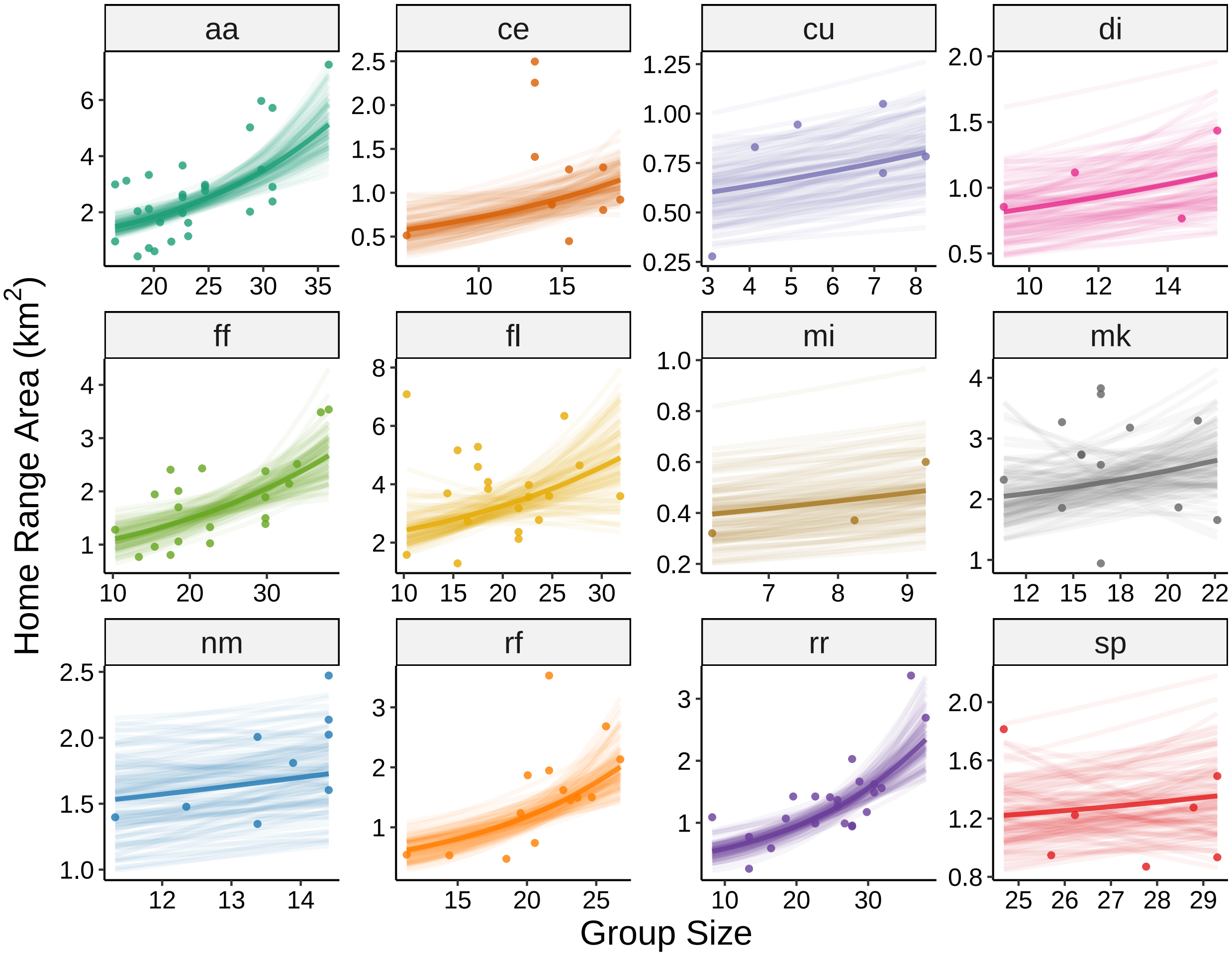


Figure S15: **Group-level model predictions for changes in home range area (HRA) as a function of group size.** Colors correspond to group identity. Lighter lines are 100 randomly sampled posterior predictions, with the posterior median as a dark line. Model was fit to $n = 156$ annual home ranges across 12 groups.

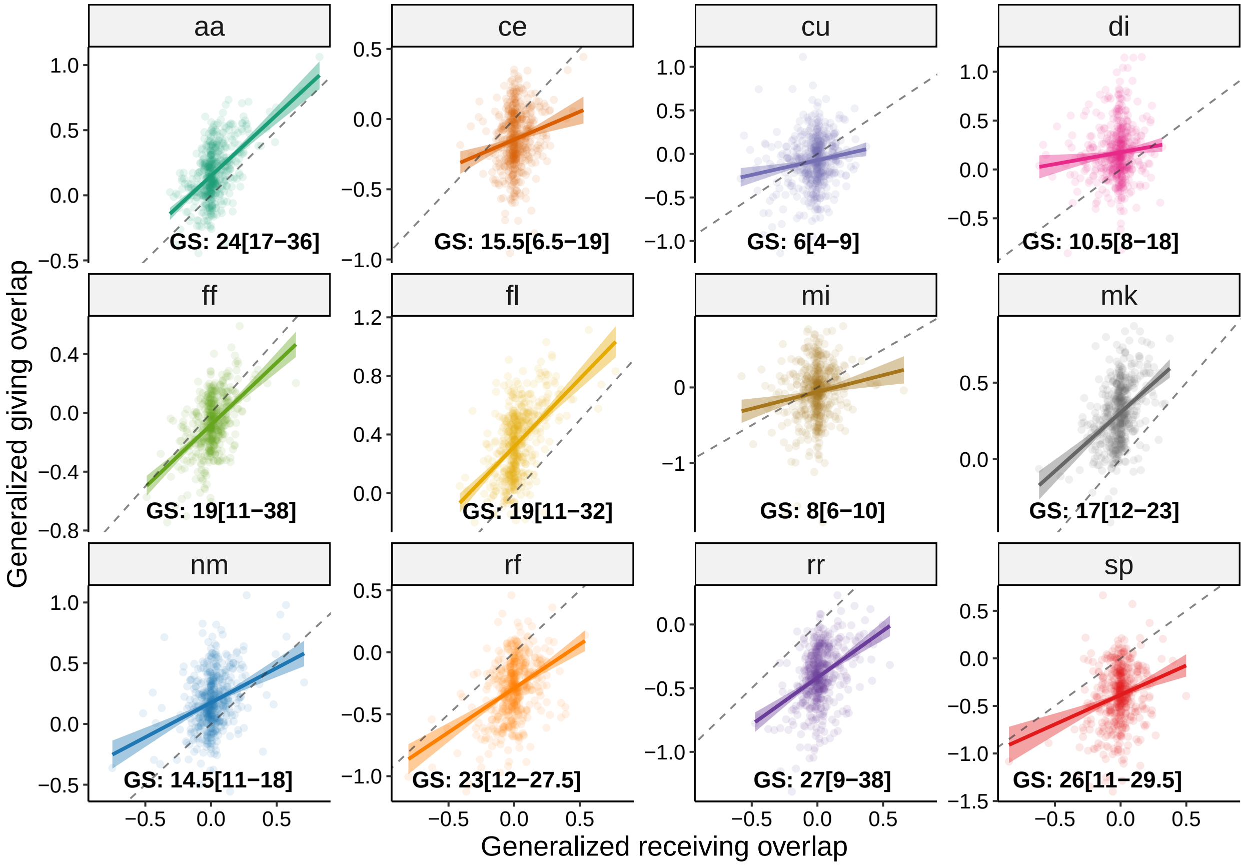


Figure S16: **Relationship between giving and receiving proportional overlap ($\mathbf{PO}_{fn}$) across groups.** Each panel displays posterior draws of a group's varying intercept for receiving overlap (x-axis; when the group is focal) against its varying intercept for giving overlap (y-axis; when the same group is the neighbor). Each point represents a posterior sample. Linear trends are fit using ordinary least squares, with 95% confidence intervals shown as shaded bands. Colors indicate group identity. Group size summaries over the full observation period (GS: median[min–max]) are annotated within each panel. Model was fit to $n = 930$ dyad-years (59 dyads, 12 groups).

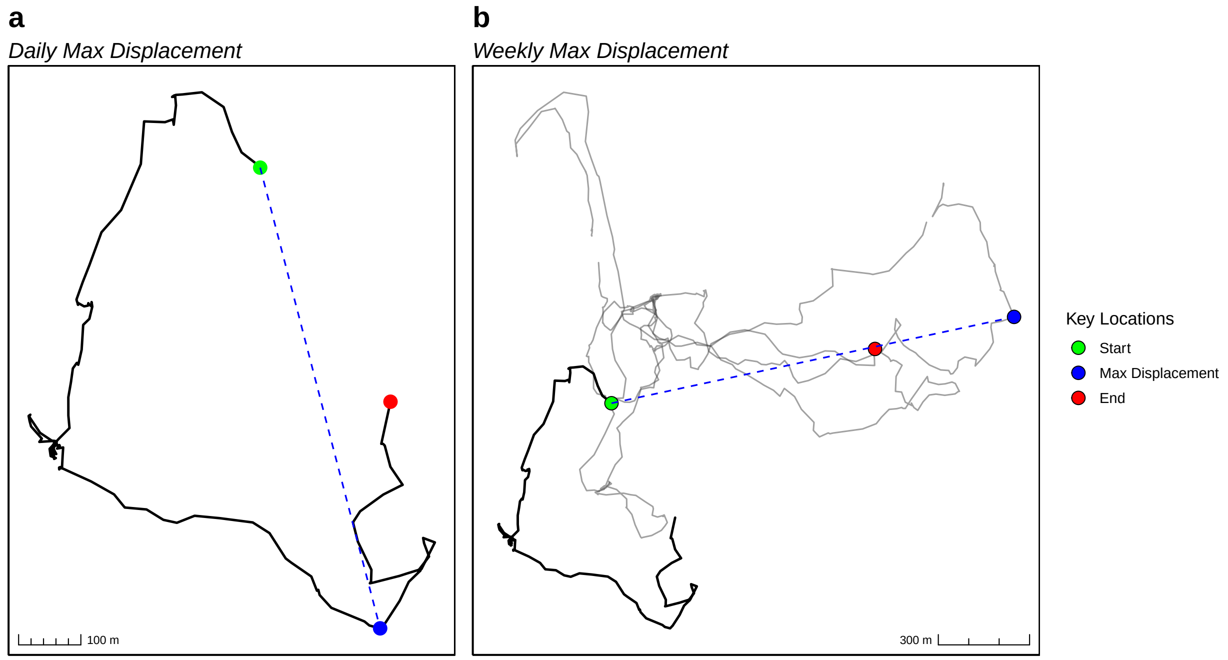


Figure S17: Visual representation of (a) daily and (b) weekly maximum displacement for a single group. In both panels, green circles mark the first recorded location, red circles mark the last location, and blue circles indicate the point of maximum displacement from the starting location. The weekly track in (b) is shown in light grey, with the corresponding daily track from (a) overlaid in a darker grey to highlight its position within the broader weekly track.

### S1.7 Temporal Scaling of Exploratory Movement

Given that larger groups had lower per capita fruit intake and larger home ranges, we expected they should also exhibit longer daily path lengths. This expectation stems from the assumption that larger groups deplete food patches more rapidly and therefore must travel farther to locate additional resources [29]. However, we found no relationship between group size and daily path length.

To investigate this apparent discrepancy, we examined whether larger groups revisit previously used areas less frequently. Our analysis indicated they do, and, together with their larger home ranges, this suggests they access a wider range of fruiting patches (with some likely being less-depleted and more nutritious) thereby reducing the need for extended daily travel.

To further explore the temporal scale of this behavior, we analyzed movement patterns using displacement metrics, which are commonly used to quantify exploration and revisitation [30, 31]. Repeated use of familiar areas is typically associated with shorter displacements over time [32]. We used maximum displacement— the farthest Euclidean distance traveled from a group's first observed location— as a proxy for exploratory behavior. This was calculated at both daily and weekly scales to capture short- and longer-term movement

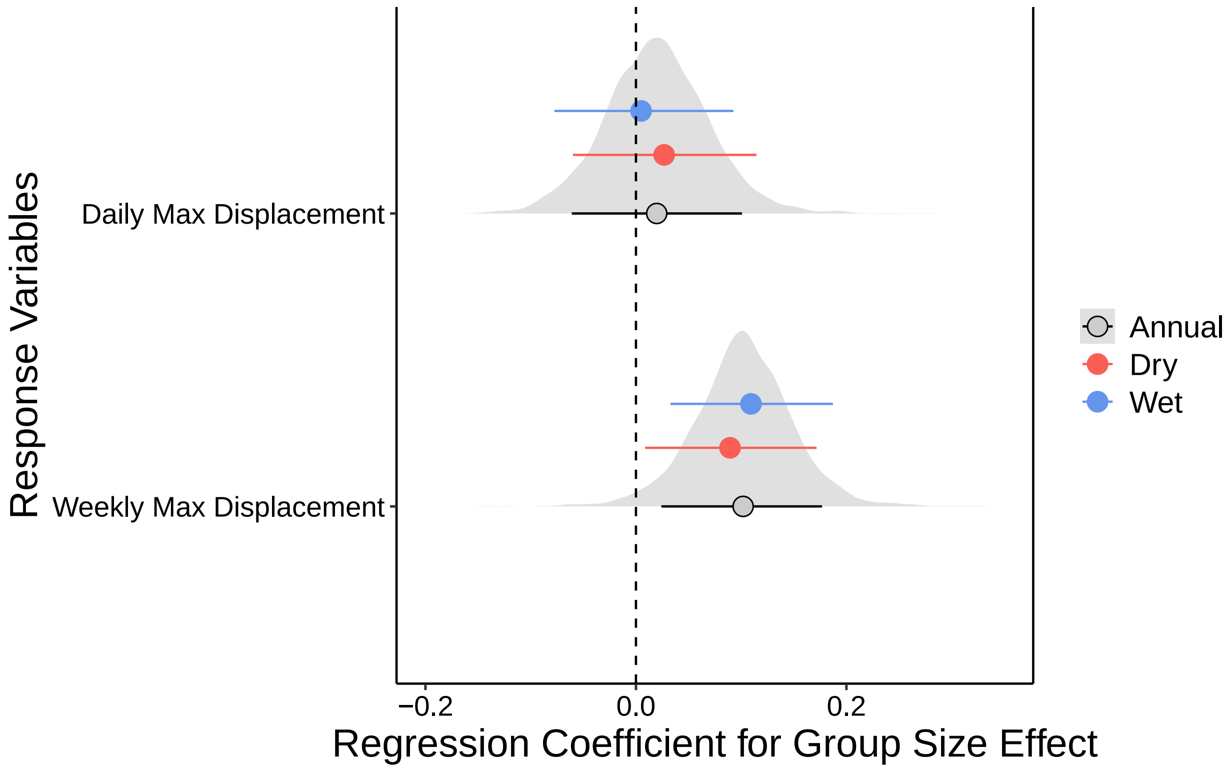


Figure S18: **Posterior estimates of the effect of group size on daily and weekly max displacement.** Points indicate median posterior estimates, and solid lines represent 89% Highest Posterior Density Intervals (HPDIs). Blue and red point intervals correspond to wet and dry season effects, respectively, while black/grey intervals indicate annual (non-seasonal) effects. Grey density slabs show the full posterior distributions for the annual effects. Daily displacement model was fit to $n = 1000$ group-days across 11 groups; weekly displacement model was fit to $n = 374$ group-weeks across 11 groups.

patterns (see Figure S17 for a visual representation).

To ensure reliable sampling, we included only days with at least 11 unique observation hours and weeks with at least 35, representing the best-sampled periods (approximately the top 30% of the dataset). We then fit Gamma-distributed GLMMs to estimate the effect of group size on both daily and weekly maximum displacement. Posterior estimates showed little to no effect of group size on daily maximum displacement, but a consistent positive association with weekly maximum displacement (Figure S18). These results indicate that larger groups do not travel farther within a single day but do expand their spatial exploration over longer time scales. This suggests that the temporal scale of movement is critical: rather than increasing daily travel, larger groups may mitigate within-group competition by gradually accessing new areas over multiple days or weeks.

### S1.8 Validation of encounter and revisitation estimates

#### Independent measures of encounters and revisitations

To evaluate whether *CTMM-derived annual encounter probabilities* correspond to observed intergroup encounter frequencies, we compared the CTMM predictions to empirical encounter counts for each dyad-year. Empirical counts were obtained from all recorded intergroup encounter locations collected between September 2009 and March 2020. These locations were recorded by observers on the ground following single groups using handheld GPS, and group identities were documented when possible. Because encounter detectability depends on how often each group was tracked, we quantified annual observation effort by counting the number of unique hours each group was observed in each year according to the GPS data-set. For each group-dyad, observation effort was quantified using the mean of the two groups' annual observation hours.

We validated *CTMM-derived revisitation rates* using an empirically derived (geometric) revisitation metric computed directly from the GPS trajectories with the `recurse` `R` package [33]. For each group-year, we generated a regular grid of points spaced 50 m apart within the group's corresponding annual 95% AKDE home-range polygon (Figure S20). Revisitation to each grid cell was quantified by counting instances in which the group returned to within 50 m of a grid point after being away for at least four hours. This four-hour temporal threshold helps ensure that revisits correspond to discrete returns to an area, rather than GPS location error or short-term movement noise. For each cell, we defined the number of revisits as the number of returns beyond the first visit and summed these across all grid cells to obtain the total number of revisitation events per group-year. Because the total number of grid cells and the amount of observation time both influence the opportunity to detect revisits, we first divided total revisitation events by the number of grid cells to obtain the mean number of revisits per 50 m grid cell per group-year, and then modeled this quantity with an offset for the number of unique observation hours per year. Model-estimated revisitation rates are therefore interpretable as revisits per hour per 50 m grid cell.

#### Model-based validation of CTMM-derived rates

We fit separate Bayesian models to relate CTMM-derived rates to their corresponding empirical measures. For encounters, we fit a zero-inflated negative binomial model in which the number of inter-group encounters observed per dyad in a year was predicted by standardized CTMM-derived encounter probability, with a varying intercept for year and observation effort as an offset to account for unequal tracking effort. For revisitation, we fit a hurdle-gamma model in which this empirically-derived revisitation rate was predicted by standardized CTMM-derived revisitation rate, including an offset for annual observation hours.

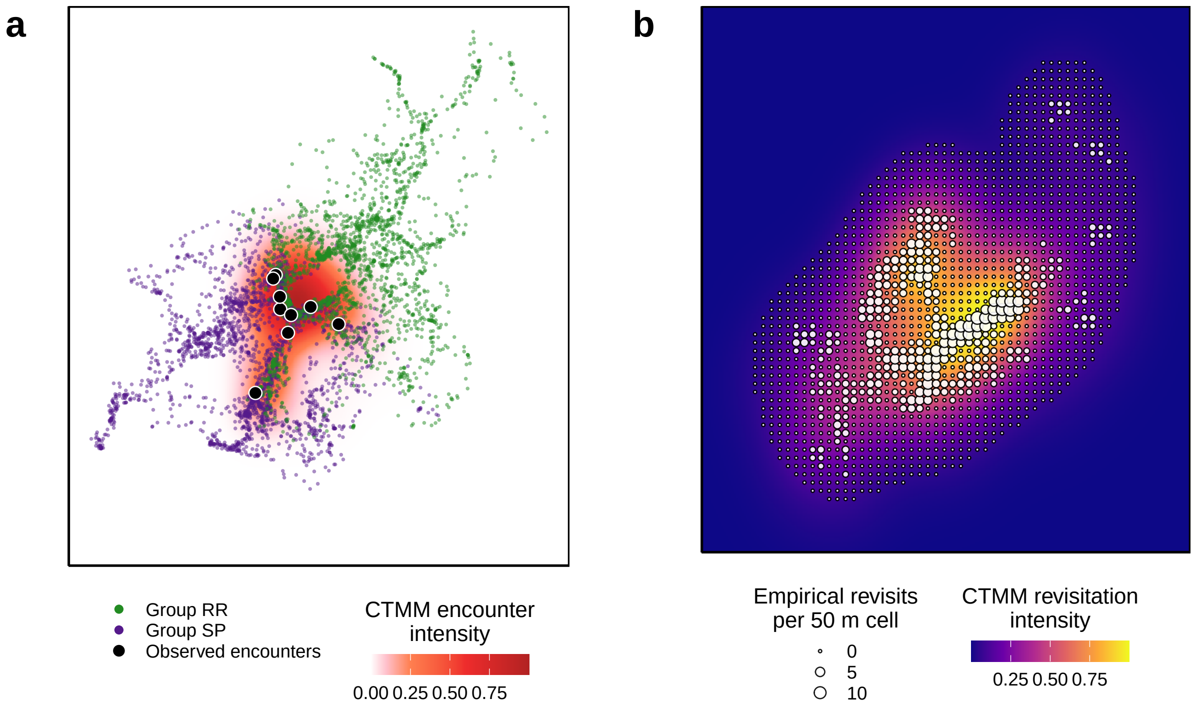


Figure S19: **Visual examples of encounter and revisitation validation.** (a) Conditional encounter distribution (CDE) between two groups (SP and RR) in 2010, shown as a CTMM-derived encounter intensity surface (warmer colors = higher predicted encounter probability). Small colored points show observed GPS location data for each group, and large black points show observed intergroup encounters between them. (b) Example of revisitation estimation for a single group in 2017. The background surface shows the CTMM-derived revisitation intensity, while overlaid points represent empirical revisits to 50-m grid cells based on the trajectory-based `recurse` approach (larger points = more revisits).

To evaluate how well CTMM-derived rates align with empirical observations, we used both continuous comparisons and binned *calibration plots*. In the binned approach, CTMM-derived values were divided into quartiles spanning low to high predicted rates, and empirical responses were averaged within each bin. This procedure is analogous to reliability or calibration diagnostics in predictive modeling: if CTMM-derived rates are well calibrated, empirical mean values should increase monotonically across bins and match the model's posterior-predicted expectations. Binning also provides a robust summary that is less sensitive to zero inflation and overdispersion in the raw responses. Empirical uncertainty was quantified using bootstrap resampling to derive 89% intervals, which capture sampling variability without parametric assumptions and align with the models' 89% HPDIs. We additionally visualized the continuous relationships by overlaying raw empirical values with each model's fitted mean trend and uncertainty.

**CTMM-derived rates show strong calibration and agreement**

Posterior estimates from the encounter model showed strong support for a positive relationship between CTMM encounter probability and observed encounter counts ($\beta = 0.79[0.64, 0.93]$; $PP > 0 = 1.00$), indicating that dyads with higher predicted encounter probability experienced more observed encounters. Similarly, posterior estimates from the revisitation model showed strong support for a positive association between CTMM-derived revisitation rate and trajectory-based revisits based on the `recurse` package functionality ($\beta = 0.58[0.48, 0.69]$; $PP > 0 = 1.00$). Both binned and continuous visualizations revealed approximately monotonic increases in encounter frequency and revisitation intensity with higher CTMM-derived estimates (Figure S20). While the empirical measures and CTMM-derived rates show broad agreement, CTMM-based estimates remain preferable because raw revisitation and encounter counts are strongly affected by sampling effort and non-random detection bias. For instance, intergroup encounters are sometimes harder for observers to detect when following larger, more dominant groups: adult males frequently move ahead of or away from the core group to engage in active aggression or displays with out-group males, placing the actual encounter outside the observers' auditory or visual range. By contrast, when observers are following smaller groups, intergroup encounters are usually obvious, as most group members flee together. CTMM estimates overcome these detection asymmetries by inferring encounters and reuse from movement data and explicitly incorporate uncertainty linked to sampling effort, also providing estimates that are less dependent on direct detectability.

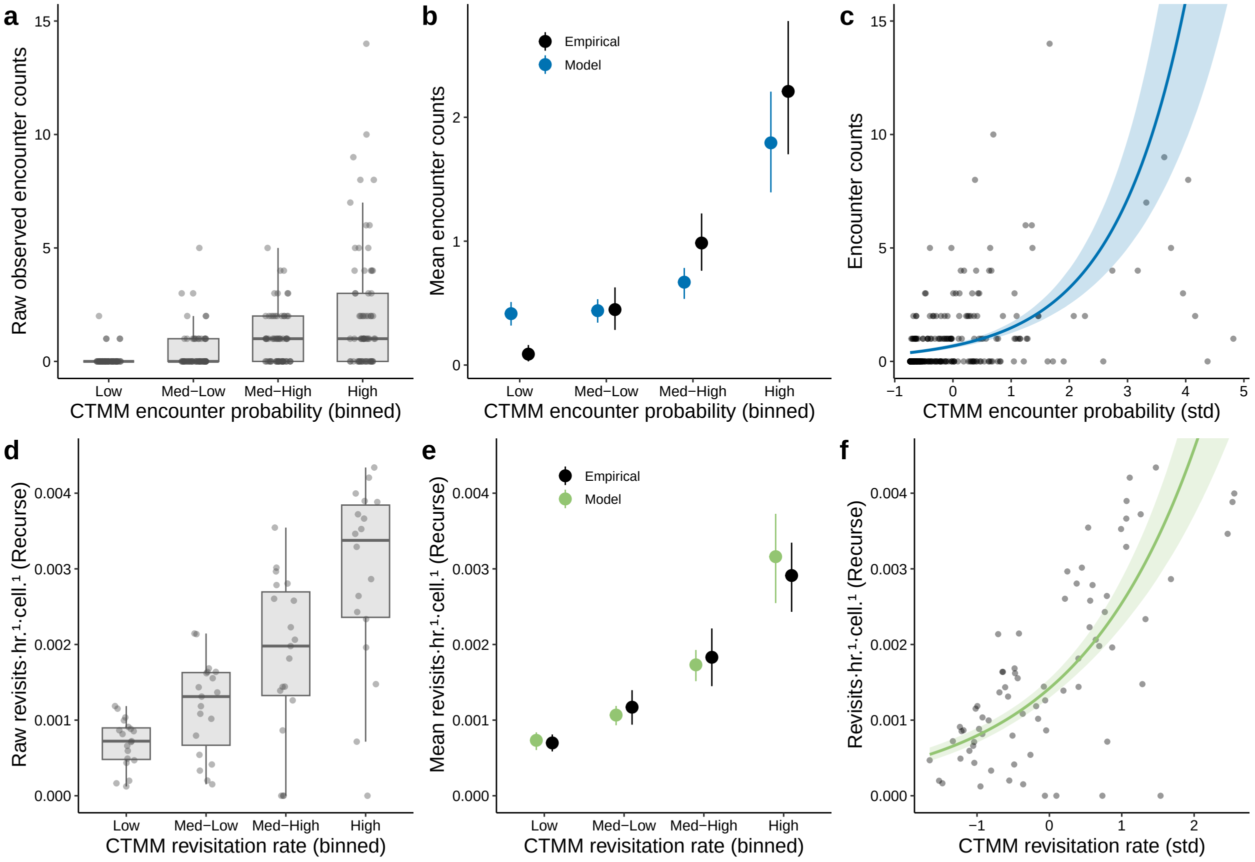


Figure S20: **Validation of CTMM-derived encounter and revisitation rates.** Panels a–c show validation for *encounter rates* (dyad-year level), and panels d–f show validation for *revisitation rates* (group-year level). Across both metrics, the left column (a,d) displays raw observations binned into quartiles of standardized CTMM-derived rates; the middle column (b,e) compares empirical bin means (black points with 89% bootstrap intervals) to posterior expected means from Bayesian models (colored points with 89% HPDIs); and the right column (c,f) shows continuous model-predicted relationships with 89% credible intervals. Encounter model predictions are shown in blue and revisitation predictions in green. Raw encounter values represent the number of intergroup encounters observed per dyad-year. Raw revisitation values represent revisits per hour per 50 m grid cell, derived from GPS trajectories by counting returns to 50 m grid cells within each group's 95% annual home range and scaled by the number of cells. Model predictions for both encounter and revisitation rates account for variation in observation effort via offsets for annual observation hours. Revisitation rate validation model was fit to $n = 101$ group-years (11 groups); encounter rate validation model was fit to $n = 269$ dyad-years (42 dyads, 11 groups).
